# Cross-tissue, single-cell stromal atlas identifies shared pathological fibroblast phenotypes in four chronic inflammatory diseases

**DOI:** 10.1101/2021.01.11.426253

**Authors:** Ilya Korsunsky, Kevin Wei, Mathilde Pohin, Edy Y. Kim, Francesca Barone, Joyce B. Kang, Matthias Friedrich, Jason Turner, Saba Nayar, Benjamin A. Fisher, Karim Raza, Jennifer L. Marshall, Adam P. Croft, Lynette M. Sholl, Marina Vivero, Ivan O. Rosas, Simon J. Bowman, Mark Coles, Andreas P. Frei, Kara Lassen, Andrew Filer, Fiona Powrie, Christopher D. Buckley, Michael B. Brenner, Soumya Raychaudhuri

## Abstract

Pro-inflammatory fibroblasts are critical to pathogenesis in rheumatoid arthritis, inflammatory bowel disease, interstitial lung disease, and Sjögren’s syndrome, and represent a novel therapeutic target for chronic inflammatory disease. However, the heterogeneity of fibroblast phenotypes, exacerbated by the lack of a common cross-tissue taxonomy, has limited the understanding of which pathways are shared by multiple diseases. To investigate, we profiled patient-derived fibroblasts from inflamed and non-inflamed synovium, intestine, lung, and salivary glands with single-cell RNA-sequencing. We integrated all fibroblasts into a multi-tissue atlas to characterize shared and tissue-specific phenotypes. Two shared clusters, CXCL10^+^CCL19^+^ immune-interacting and SPARC^+^COL3A1^+^ vascular-interacting fibroblasts were expanded in all inflamed tissues and additionally mapped to dermal analogues in a public atopic dermatitis atlas. We further confirmed these human pro-inflammatory fibroblasts in animal models of lung, joint, and intestinal inflammation. This work represents the first cross-tissue, single-cell fibroblast atlas revealing shared pathogenic activation states across four chronic inflammatory diseases.

## Introduction

Fibroblasts are present in all tissues and adopt specialized phenotypes and activation states to perform both essential functions in development, wound-healing, and maintenance of tissue architecture, as well as pathological functions such as tissue inflammation, fibrosis, and cancer responses (Koliaraki et al., 2020). Recent studies of chronic inflammatory disease have leveraged advances in high-throughput single-cell genomics, particularly single-cell RNA-sequencing (scRNAseq) to identify molecularly distinct fibroblast populations associated with pathological inflammation in different anatomical sites (Adams et al., 2020; Habermann et al., 2020; Huang et al., 2019; Kinchen et al., 2018; Martin et al., 2019; Mizoguchi et al., 2018; Smillie et al., 2019; Zhang et al., 2019). A study of the large intestine from patients with ulcerative colitis (UC) identified stromal cells expressing Oncostatin-M receptor (OSMR) enriched in biopsies tracking with failure to respond to anti-TNF therapy (West et al., 2017). Further studies suggested immunomodulatory roles for OSMR^+^ intestinal fibroblasts through interactions with inflammatory monocytes (Smillie et al., 2019) and neutrophils (Friedrich et al., 2020). Lung investigations identified that COL3A1^+^ACTA2^+^ myofibroblasts, PLIN2^+^ lipofibroblast-like cells, and FBN1^+^HAS1^+^ fibroblasts are expanded in lung biopsies from patients with idiopathic pulmonary fibrosis (IPF) (Adams et al., 2020; Habermann et al., 2020). In the salivary gland, chronic destructive inflammation in primary Sjögren’s syndrome (pSS) with tertiary lymphoid structures is linked to the expansion of PDPN^+^CD34^−^ fibroblasts (Nayar et al., 2019). In the synovial tissue, FAP*α*^+^CD90^+^ fibroblasts are expanded in patients with rheumatoid arthritis (RA) (Wei et al., 2020; Zhang et al., 2019) and drive leukocyte recruitment and activation in an animal model of arthritis (Croft et al., 2019).

In each study, inflammation-associated fibroblasts are characterized by their ability to produce and respond to inflammatory cytokines. These cytokines are often members of conserved families that signal through similar downstream pathways and result in similar effector functions (West, 2019). For instance, the inflammatory cytokines IL-6, Oncostatin M (OSM), leukemia inhibitory factor (LIF), and IL-11 all belong to the gp130 family, whose cognate receptor molecules, including IL-6R, OSMR, LIFR, and IL-11R, contain the Glycoprotein 130 (gp130) subunit. In UC, OSMR^+^ fibroblasts express high levels of the IL-11 encoding gene (Smillie et al., 2019). In RA, a subset of FAP*α*^+^CD90^+^ synovial fibroblasts produce high levels of IL-6 (Zhang et al., 2019) through an autocrine loop involving LIF and LIFR (Nguyen et al., 2017; Slowikowski et al., 2019). In a mouse model for human IPF, IL-11 producing fibroblasts drive both fibrosis and chronic pulmonary inflammation (Ng et al., 2020). These examples of gp130-family cytokines associated with pro-inflammatory fibroblasts highlight that while individual factors may be tissue-specific, their downstream effects may be shared across diseases. This pattern underlines an important question with clinical implications: are inflammation-associated fibroblasts tissue-specific or do they represent shared activation states that manifest a common phenotype across different diseases? A drug that targets a shared pathogenic phenotype can potentially be used to treat multiple inflammatory diseases. Identifying such shared fibroblast programs presents a major challenge, as these programs are likely to be transient and reversible activation states that vary over the course of a disease, rather than representing a static, committed cell lineage (Wei et al., 2020).

The identification of shared cell states across tissues with scRNAseq has recently become possible with advances in statistical methods for integrative clustering (Butler et al., 2018; Korsunsky et al., 2019; Tran et al., 2020) and reference mapping (Andreatta et al., 2020; Kang et al., 2020; Lotfollahi et al., 2020). Integrative clustering identifies similar cell states across a range of scRNAseq datasets, even when the datasets come from different donors, species, or tissues. For example, using integrative clustering, Zhang et al., 2020 identified shared macrophage activation states across five tissues, and Butler et al., 2018 identified shared pancreatic islet cells between mouse and human datasets. Reference mapping allows rapid comparison of data from a new study to a well annotated reference, even if the study represents a tissue, disease, or species not present in the reference atlas. For instance, Andreatta et al., 2020 mapped T cell subtypes to a scRNAseq atlas of annotated tumor infiltrating T cells, while Lotfollahi et al., 2020 found disease-related immune states by mapping PBMCs from patients with COVID19 to a healthy reference library of immune cells.

In this study, we generated single-cell RNAseq profiles of patient-derived CD45^−^ stromal cells and then characterized fibroblasts across multiple inflammatory diseases involving lung, intestine, salivary gland and synovium. After confirming known fibroblast subtypes in our data, we built a *de novo*, integrated fibroblast atlas and identified five shared phenotypes, two of which are consistently expanded in all four inflammatory diseases. Using reference mapping, we map these to human dermal fibroblasts from inflamed and healthy skin and to fibroblasts from mouse models of lung, synovial, and intestinal inflammation to demonstrate the generalizability of our findings. Our integrated resource represents the first systematic examination of fibroblast subsets and activation states in inflamed tissues. Our identification of two pathogenic fibroblast phenotypes that are shared amongst four inflammatory diseases novel avenues for therapeutic targeting. By making available the necessary computational tools to map new datasets to our annotated fibroblast atlas, we provide a common reference for future studies of fibroblasts in tissues and diseases.

## Results

### Single-cell transcriptional profiles of fibroblasts in human lung, salivary gland, synovium, and intestine

We used droplet-based scRNAseq to profile individual fibroblasts from a total of 74 high quality samples in lung, large intestine, lip salivary glands, and joint synovium, selecting donors with inflammatory diseases and controls (**Figure 1a**). In synovium, we collected arthroplasties and biopsies from 18 patients with RA and 6 with osteoarthritis (OA) (**Supplementary Table 1**). In the intestine, we collected large intestinal biopsies from patients with UC (n=8) and control (n=5) donors (**Supplementary Table 2**). Included in the 8 UC samples were 4 patients for whom we had paired inflamed and adjacent non-inflamed tissue biopsies. For the lung analysis, we acquired lung tissue samples from 19 patients with ILD and 4 control samples from donor lungs (**Supplementary Table 3**). To examine salivary glands, we used lip biopsy tissue from 7 patients with primary Sjögren’s Syndrome (pSS) and 6 patients with a non-Sjögren’s Sicca syndrome, characterized as non-autoimmune dryness, as control-comparators (**Supplementary Table 4**). In order to enrich for stromal cells, we used flow cytometry to sort live, CD45^−^EpCAM^−^ cells from intestine and synovium samples (**Figure 1a**), depleting CD45^+^ immune and EpCAM^+^ epithelial populations (**Supplementary Figure 1a**). We avoided this strategy in the salivary gland, in order to optimize cell numbers in small biopsies, and in the lung, in which flow cytometry compromised fibroblast cell yields. We performed droplet-based scRNAseq (10x Genomics) on all samples, applied stringent QC to remove low quality libraries and cells (**Supplementary Figure 1b-d**), and combined all data samples to analyze 221,296 high quality cells. Using clustering analysis (**Methods**), we identified 7 major cell types (**Figure 1b**) with canonical markers (**Figure 1c**): *CDH5*^+^ endothelial cells, *COL1A1*^+^ fibroblasts, *EPCAM*^+^ epithelial cells, *GFRA3*^+^ glial cells, *JCHAIN*^+^ plasma cells, *MCAM*^+^ perivascular murals, and *PTPRC*^+^ leukocytes. Consistent with our flow sorting strategy, non-stromal cells (epithelial, glial, and immune) were more abundant in the salivary gland and lung (**Supplementary Figure 1e**). Importantly, we identified stromal (endothelial, mural, and fibroblast) populations in all four tissues, allowing us to carry out a focused analysis of fibroblasts across tissues.

**Figure 1.**
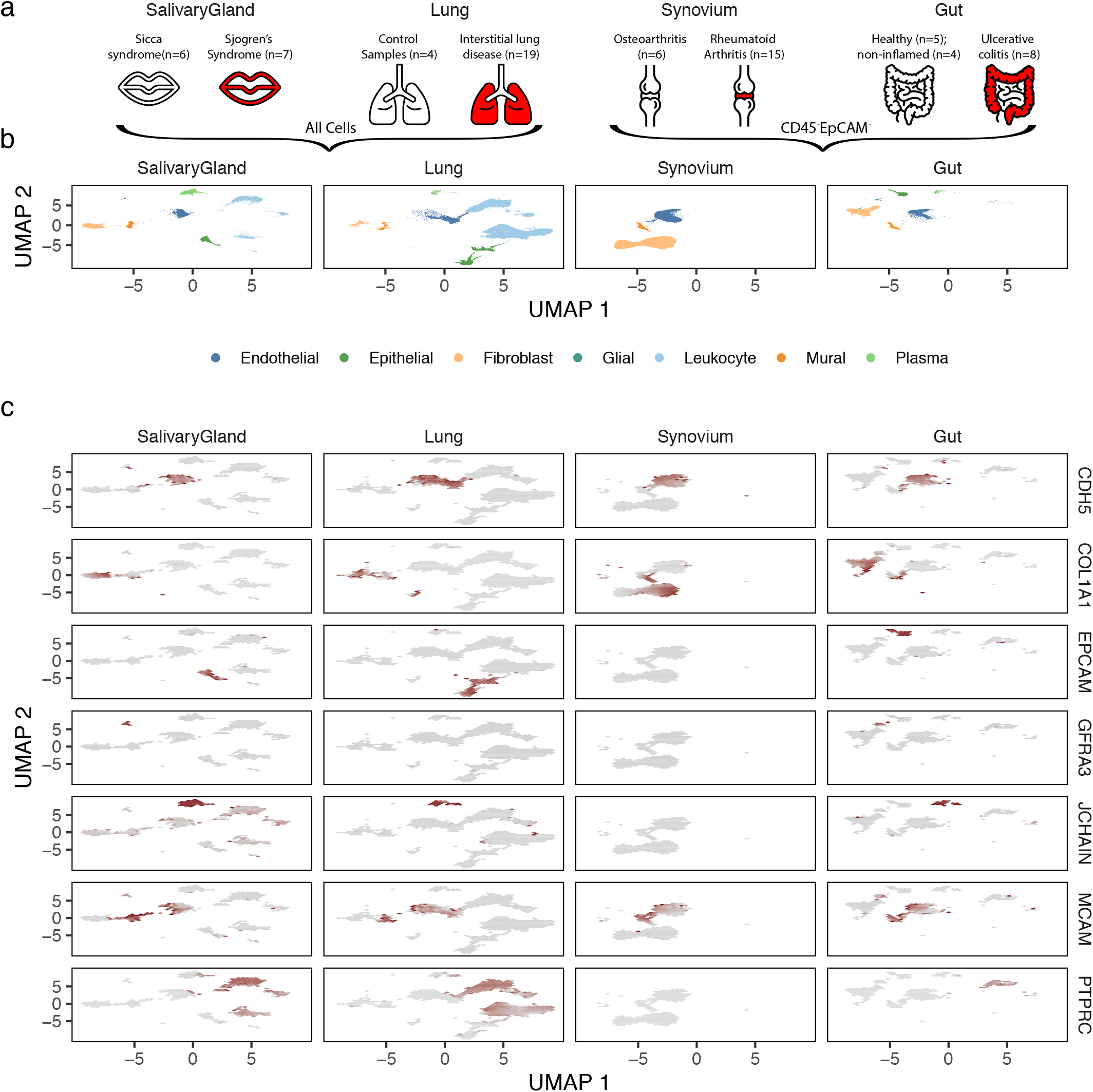
scRNAseq profiles of intestine, lung, salivary gland, and synovium. (a) Surgical samples were collected from intestine, lung, salivary gland, and synovium, from patients with inflammatory-disease and appropriate controls. After tissue disaggregation, all cells from lung and salivary gland and CD45-EpCAM-cells from synovium and intestine were profiled with scRNAseq and (b) analyzed to identify fibroblasts and other major cell types. (c) Cell type annotation was performed with known markers for each major population.

### Fibroblast heterogeneity within tissues

We next examined the heterogeneity of fibroblast cell states within individual tissues. We performed a separate fine-grained clustering analysis for fibroblasts within each of the four tissues and annotated clusters with previously identified states (**Figure 2a**) by comparing published marker genes (**Supplementary Figure 2a-d**) with cluster markers in our data (**Supplementary Table 5**). In the intestine, we were able to recapitulate 7 of 8 populations identified in (Smillie et al., 2019): crypt-associated WNT2B^+^Fos^hi^ and WNT2B^+^Fos^lo^, epithelial-supportive WNT5B^+^-1 and WNT5B^+^-2, stem cell niche supporting RSPO3^+^, inflammatory, and myofibroblasts. We note that our data did not support the 2 subtypes of WNT2B^+^Fos^lo^ fibroblasts identified originally in (Smillie et al., 2019). In the lung, Habermann et al., 2020 described 4 states: HAS1^+^, PLIN2^+^, fibroblasts, and myofibroblasts. However, in their analysis, HAS1^+^ cells were identified in only 1 of 30 donors. When we re-analyzed their data to identify clusters shared by multiple donors, we could not distinguish the HAS1^+^ from PLIN2^+^ population and thus merged these two in our annotation. In the salivary gland, the only single-cell study of fibroblasts to date was performed with multi-channel flow cytometry (Nayar et al., 2019), not scRNAseq. The findings here represent the first set of scRNA-seq data in this context. In our single-cell clusters, we identified the two populations previously described (CD34^+^ and CCL19^+^) and confirmed the expression of key distinguishing cytokines and morphogens that they measured by qPCR (**Supplementary Figure 2b**). In the synovium, we clustered 55,143 fibroblasts into 5 major states described in three scRNAseq studies (Croft et al., 2019; Mizoguchi et al., 2018; Zhang et al., 2019). These states are largely correlated with anatomical position: THY1^−^PRG4^+^ cells in the synovial boundary lining layer and THY1^+^, DKK3^+^, HLA-DRA^+^, and CD34^+^ cells within the sublining. In total, we labeled 17 fibroblast clusters defined across all four individual tissues.

**Figure 2.**
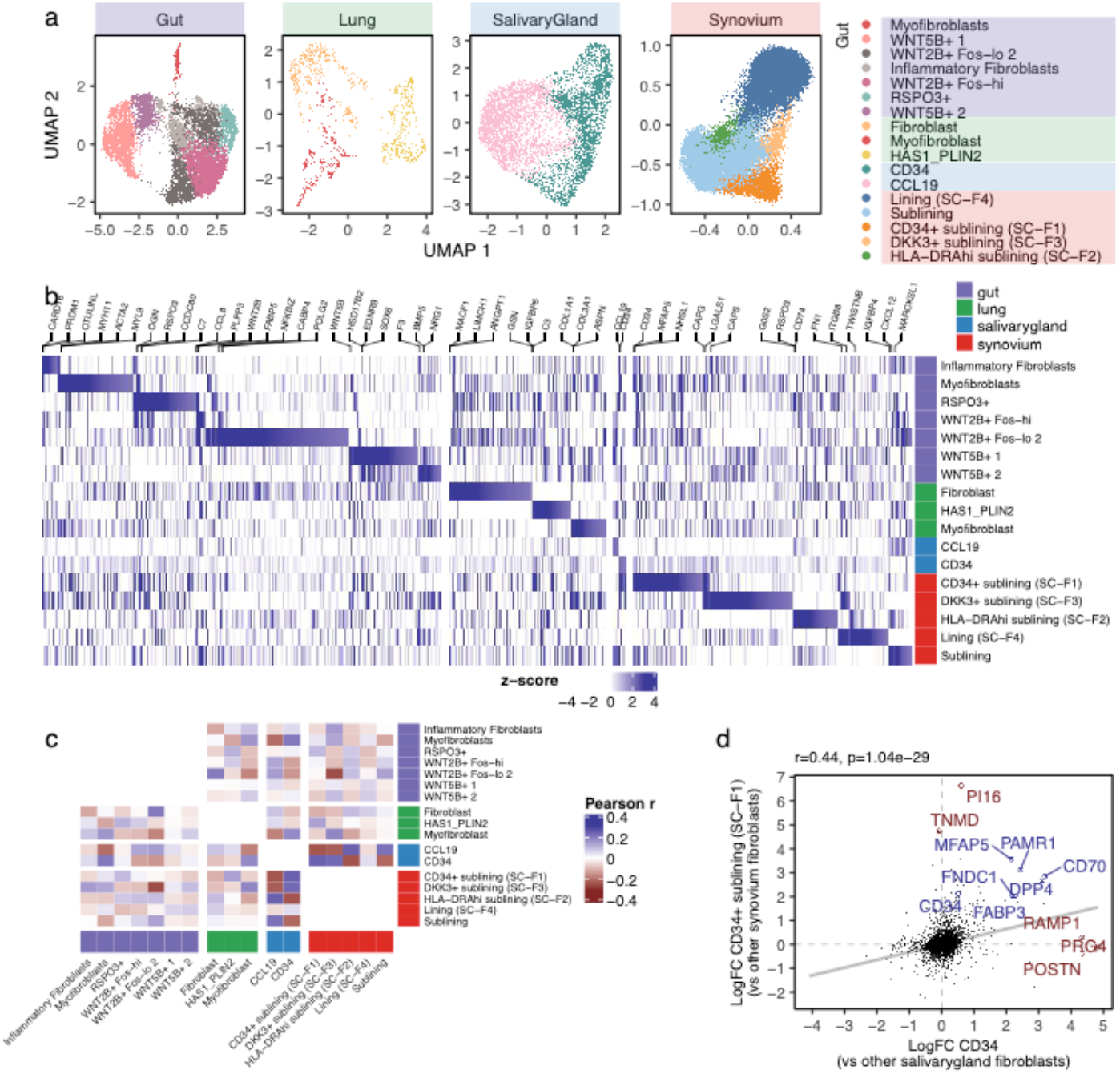
Fibroblast heterogeneity within tissues. (a) We analyzed fibroblasts separately from each tissue to identify tissue-specific subsets described in previous single-cell studies. Each panel shows a UMAP representation of fibroblasts from one tissue, labeled with clustering and marker analysis. (b) All (n=7,380) genes nominally upregulated in any cluster were plotted in a heatmap. Color denotes the log fold change, normalized by estimated standard deviation, of a gene in a cluster (versus other clusters in that tissue). Top genes for each cluster were named above the heatmap. Each row denotes a fibroblast cluster, colored by the tissue in which it was identified. (c) To compare the expression profiles of clusters across tissues, we correlated the expression values from (b) for all pairs of clusters. Here, color denotes Pearson’s correlation coefficient. (d) One highly correlated pair of clusters from salivary gland (x-axis) and synovium (y-axis) represented by scatter plots of (differential) gene expression. Blue genes are shared by the two clusters, while red genes are unique to one cluster.

Next, we asked whether fibroblast states defined within one tissue shared similar expression profiles with states defined in other tissues. We performed cluster marker analysis within each tissue, quantifying the overexpression of each gene in each cluster in terms of the log_2_ fold change with other clusters. We plotted 4,897 genes that were overexpressed in at least one cluster and labeled the top 3 markers per cluster (**Figure 2b**). We noticed that many marker genes were present in clusters from different tissues. To find which pairs of clusters across tissues were most similar, we correlated (differential) expression profiles (**Methods**) for cross-tissue clusters (**Figure 2c**). The most correlated (Pearson *r* = 0.44, *p* = 10^−29^) pair of clusters contained CD34^+^ fibroblasts in the salivary gland and CD34^+^ sublining (SC-F1) fibroblasts in the synovium (**Figure 2d**). Although they shared multiple marker genes (*PAMR1*, *MFAP5*, *CD34*, *CD70*, *DPP4*, *FABP3*, and *FNDC1*), they also had tissue-related, cluster-specific genes (*POSTN, RAMP1, PRG4, PI16,* and *TNMD*). The shared markers suggest a shared function. The cluster-specific genes may have arisen from a technical artefact, such as different clustering parameters in the tissue-specific analyses, or from true biological signal, such as a tissue-specific microenvironment. In order to distinguish between the two possibilities, we decided to perform a single integrative clustering analysis with fibroblasts from all tissues.

### Integrative clustering of fibroblast across tissues

To construct a cross-tissue taxonomy of fibroblast states, we pooled 55,143 synovial, 15,089 intestinal, 7,474 salivary gland, and 1,442 pulmonary fibroblasts together and performed integrative clustering analysis. The different numbers of fibroblasts from each tissue, arising from the fact that we enriched for stromal cells in intestine and synovium but not in lung and salivary gland, presented a technical challenge. The results of many analyses, including PCA, are biased towards tissues with more cells, rather than treating each tissue equally. The second major analytical challenge arises from the fact that gene expression depends on a complex interplay of tissue, donor, and cell state. As we have described in previous work (Korsunsky et al., 2019), such confounding variation is particularly challenging to model in scRNAseq data, as the confounder can have both global and cell-type specific effects on gene expression.

We designed an analytical pipeline for integrative clustering to address the two concerns described above (**Figure 3a**). In this pipeline, we select genes that were informative in the tissue-specific analyses (**Methods**), associated with either cluster identity (**Supplementary Table 5**, n=7,123) or inflammatory status (**Supplementary Table 6**, n=6,476) within tissue, for a total of 9,521 unique genes. To minimize the impact of different cell numbers, we performed weighted PCA analysis, giving less weight to cells from over-represented tissues (e.g. synovium) and more to cells from under-represented tissues (e.g. lung), such that the sum of weights from each tissue is equivalent (**Methods**). Compared to unweighted PCA, this approach results in principal components whose variation is more evenly distributed among tissues (**Supplementary Figure 3a**). As expected, in this PCA space, cells group largely by donor and tissue (**Supplementary Figure 3b,c**). In order to appropriately align cell types, we removed the effect of donor and tissue from the cells’ PCA embedding coordinates with a novel, weighted implementation of the Harmony algorithm that we developed for this specific application (**Methods**). UMAP visualization of the harmonized embeddings shows that cells from different tissues are well mixed (**Figure 3b**). In contrast, fibroblast states identified in tissue-specific analyses are well separated (**Supplementary Figure 3d**), suggesting that the integrated embedding faithfully preserves cellular composition. In this integrated space, we performed standard graph-based clustering to partition the cells into 14 fibroblast states (**Figure 3c**) with representation from all 4 tissues (**Supplementary Figure 3e**). These 14 integrated clusters represent putative shared fibroblast states, each of which may be driven by a combination of both shared and tissue-specific gene programs.

**Figure 3.**
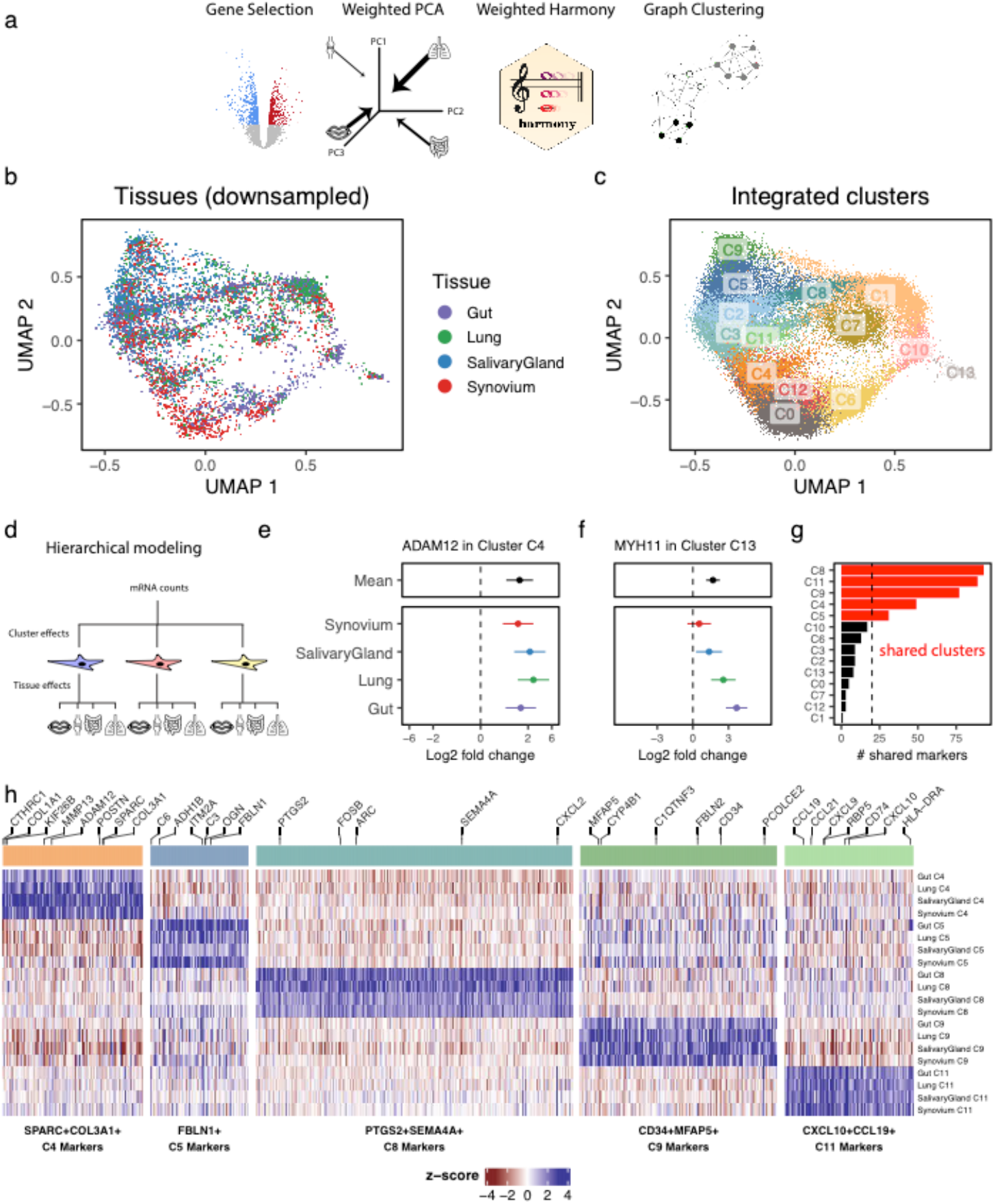
Integrative clustering and differential expression across tissues. (a) We developed a pipeline to integrate samples from multiple donors and multiple tissues with unbalanced cell numbers. The pipeline starts with gene selection, pooling together genes that were informative in single-tissue analyses. With these genes, we performed weighted PCA, reweighting cells to computationally account for the unbalanced dataset sizes among the tissues. These PCs are adjusted with a novel formulation of the Harmony integration algorithm and used to perform graph-based clustering. We applied this pipeline to all fibroblasts across tissues. (b) The integrated UMAP projection shows cells from all tissues mixed in one space. For clarity, we down-sampled each tissue to the smallest tissue, the lung, choosing 1,442 random fibroblasts from intestine, synovium, and salivary gland. (c) Graphed-based clustering proposed 14 fibroblast clusters in the integrated embedding. (d) Gene-level analysis to find upregulated marker genes for clusters was done with hierarchical regression, to model complex interactions between clusters and tissues. This strategy distinguishes cluster marker genes that are (e) tissue-specific, such as MYH11 in C13, from those that are (f) shared among tissues, such as ADAM12 in C14. Points denote log fold change (cluster vs other fibroblast) and error bars mark the 95% confidence interval for the fold change estimate. (g) The number of shared genes for each cluster, ranked from most to least, prioritizes clusters with large evidence of shared gene expression (in red) from those with little (in black). Marker genes for the 5 shared clusters plotted in a heatmap. Each block represents the (differential) gene expression of a gene (column) in a cluster, for a tissue (row).

### Identification of shared and tissue-specific marker genes in integrated clusters

Next, we modeled gene expression to define active gene programs in the 14 integrative fibroblast clusters. In particular, we wanted to distinguish between two types of cluster markers: tissue-shared and tissue-specific. Tissue-shared markers are highly expressed in the cluster for all four tissues. Tissue-specific markers are highly expressed in the cluster for at least one tissue but not highly expressed in at least one other tissue. In our expression modeling analysis, we needed to allow for the possibility that tissue gene expression will be consistent in clusters and variable in others (**Figure 3d**). As we explain in our approach below, we will use ADAM12 expression in cluster C4 as an example of a tissue-shared gene and MYH11 expression in cluster C13 as an example of a tissue-specific gene.

Typically, cluster marker analysis is done with regression, to associate gene expression with cluster identity. To address the complex interaction between cluster and tissue identity in our data, we used mixed-effects regression to perform hierarchical cluster marker analysis (**Methods**). This analysis estimated two sets of differential expression statistics for each gene: mean log_2_ fold change (e.g. cluster 0 vs all other clusters) and tissue-specific log_2_ fold change (e.g. cluster 0 in lung vs all other clusters in lung). This approach distinguishes shared marker genes, defined by minimal tissue-specific contributions, from tissue-specific marker genes, defined by large tissue-specific fold changes, relative to the mean fold change. To demonstrate, we plotted the estimated log_2_ fold changes, with a 95% confidence interval, for one shared (**Figure 3e**) and one tissue-specific (**Figure 3f**) cluster marker. ADAM12, a shared marker for cluster C4, has significant (log_2_ fold-change = 1.6, *p* = 6.5 × 10^−9^) mean differential expression in C4, while the tissue-specific effects (in color) are not significantly different for any one tissue (**Figure 3e**). In contrast, MYH11, is differentially overexpressed in cluster C13 for intestinal (log_2_ fold-change = 3.7, *p* = 8.5 × 10^−16^) and lung fibroblasts (log_2_ fold-change = 2.6, *p* = 5.9 × 10^−7^) but not for synovial or salivary gland cells (**Figure 3f**). Because MYH11 is so strongly overexpressed in intestinal and lung fibroblasts, the mean log_2_ fold-change is also significant (log_2_ fold-change = 1.7, *p* = 5.7 × 10^−9^) and therefore is not a good metric alone to determine whether a marker is shared or tissue-specific.

We defined tissue-shared cluster markers conservatively by requiring a marker gene to be significantly overexpressed in all four tissues, such as *ADAM12* above. With this criterion, we quantified the number of shared marker genes per cluster (**Figure 3g**). Clusters C0, C1, C2, C3, C6, C7, C10, C12, and C13 each had fewer than 20 shared markers. Based on this cutoff, we decided that these clusters had too little evidence of shared marker genes to be reliably called shared clusters. We assigned names for the remaining clusters based on their shared gene markers: SPARC^+^COL3A1^+^ C4, FBLN1^+^ C5, PTGS2^+^SEMA4A^+^ C8, CD34^+^MFAP5^+^ C9, and CXCL10^+^CCL19^+^ C11. We then plotted the log_2_ fold change values of all 1,524 shared markers for these clusters in **Figure 3h** and report the results of the full differential expression analysis in **Supplementary Table 7**.

### Identification of fibroblast states expanded in inflamed tissue

We next addressed which cross-tissue fibroblast states were expanded in inflamed tissues. In order to perform this association across tissues, we first needed to define a common measure of tissue inflammation. While histology is often the gold standard to assess inflammation, histological features are inherently biased to tissue-specific pathology. Instead, we decided to define inflammation in a tissue-agnostic way, as the relative abundance of immune cells in each sample. While immune cell abundance alone oversimplifies complex pathological processes, it is a ubiquitous and quantifiable measure of chronic inflammation. We quantified the fraction of immune cells based on previously labeled scRNAseq clusters (**Figure 1b**), for salivary gland and lung samples, and based on the proportion of CD45^+^ cells by flow cytometry (**Supplementary Figure 1a**), for synovium and intestine (**Figure 4a**). We note that these estimates are quantified with dissociated cells from cryopreserved tissue (**Methods**) and thus lack granulocytes, such as neutrophils, which constitute an important part of tissue inflammation. In order to get comparable results across tissues, we standardized the raw tissue-specific immune cell frequencies to a common scale from 0 (not inflamed) to 1 (inflamed) (**Figure 4b**). Importantly, this transformation (**Methods**) removes the impact of distributional differences among tissues and preserves the order of scores within each tissue.

**Figure 4.**
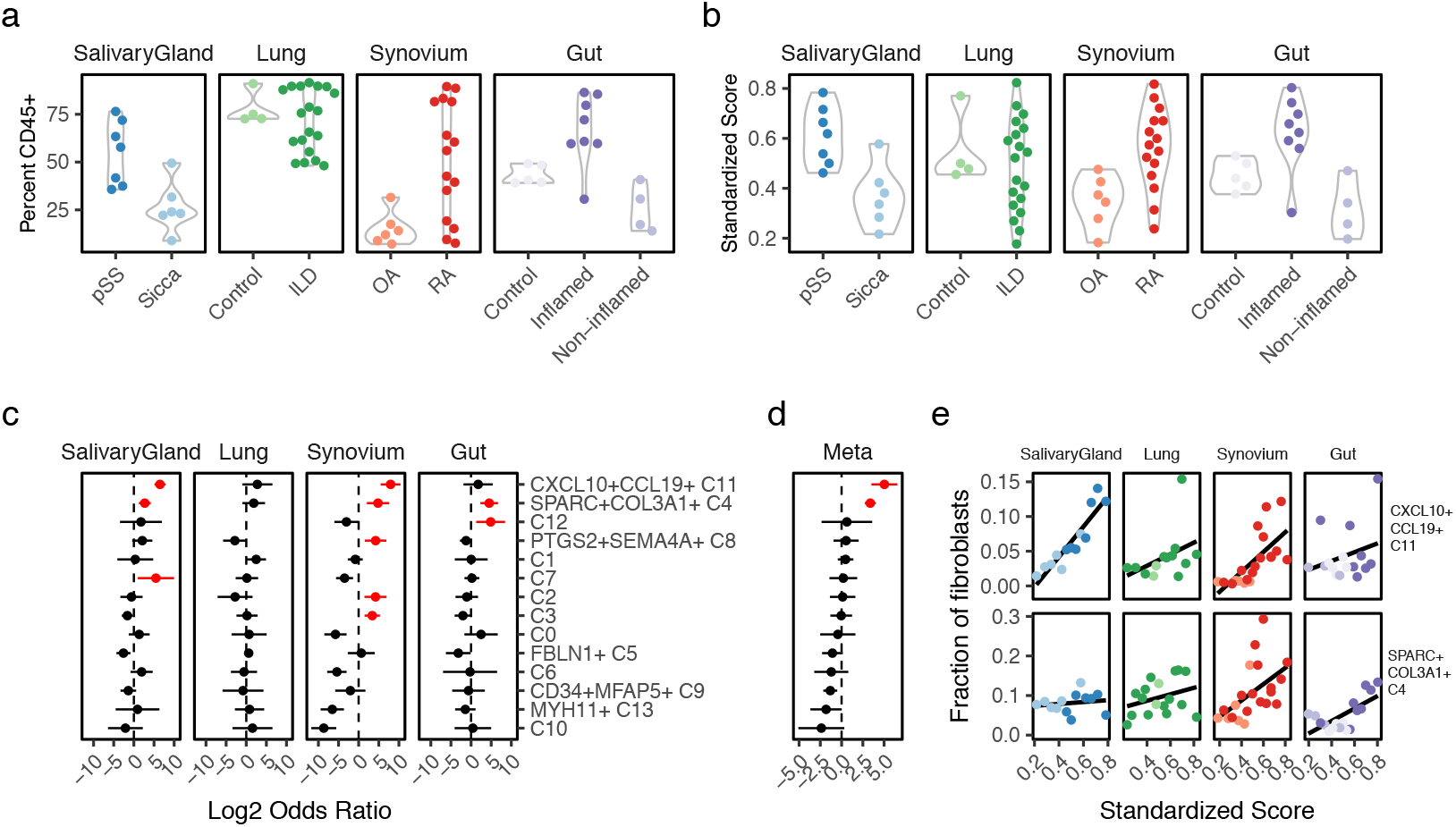
Sample level inflammation scores. We computed the relative abundance of CD45^+^ immune cells to all cells in each sample. (b) We standardized these frequencies across tissues into an inflammation score that ranges from 0 to 1 and removes distributional differences. (c) Association analysis results between fibroblast cluster abundance and standardize inflammation scores. Here, each point represents the log fold change in fibroblast cluster abundance with increasing inflammation and the line represents that point’s 95% confidence interval. Red denotes estimates with one-tailed FDR<5%. (d) The tissue specific results were summarized using meta-analysis. (e) For CXCL10+CCL19+ (C11) and SPARC+COL3A1+ (C4) fibroblasts, scatterplots relating to standardized inflammation scores (x-axis) to relative fibroblast frequency (y-axis).

Using these standardized inflammation scores, we performed a separate association analysis with mixed-effects logistic regression for each tissue. This analysis provided, for each tissue and fibroblast state, the effect of increased inflammation on cluster abundance (**Figure 4c**). Positive log odds ratios denote expansion with inflammation whereas negative ratios denote a diminishing population. Some clusters, such as C2, C3, C7, PTGS2^+^SEMA4A^+^ C8, and C12, were significantly (FDR<5%, red) expanded in only one tissue. Others, such as CXCL10^+^CCL19^+^ C11 and SPARC^+^COL3A1^+^ C4, were significantly expanded in multiple tissues. We confirmed that association with normalized inflammation scores did not change the qualitative results within tissue but did make the results more interpretable across tissues (**Supplementary Figure 4**). We then performed a meta-analysis of these tissue-specific effects (**Methods**) to prioritize clusters expanded consistently across all tissues (**Figure 4d**). This meta-analysis identified two fibroblast states significantly expanded in inflamed samples from all 4 tissues (**Figure 4e**): SPARC^+^COL3A1^+^ (C4) (*OR* = 10.4, 95% *CI*[6.6, 16.2], *p* = 9.4 × 10^−25^), and CXCL10^+^CCL19^+^ (C11) fibroblasts (log *OR* = 32.7, 95% *CI* [11.4, 94.0], *p* = 9.6 × 10^−11^). The reported odds ratio values denote the odds of a cell being in a cluster (versus not) given that it came from an inflamed sample. Because the effects for these clusters were similar across tissues, pooling in the meta-analysis increased the power to detect these abundance changes.

### Distinct immune-interacting and vascular-interacting fibroblast states expanded in tissue inflammation

The two fibroblast states consistently expanded in inflamed tissue are characterized by distinct gene programs (**Figure 5a**) that reflect putative distinct functions during tissue inflammation. To explore these potential roles, we performed gene set enrichment analysis with 6,369 Gene Ontology (Ashburner et al., 2000) and 50 MSigDB hallmarks pathways (Liberzon et al., 2011) (**Supplementary Table 8, Figure 5b**). Marker genes for CXCL10^+^CCL19^+^ fibroblasts were enriched for pathways involved in direct interaction with immune cells, including lymphocyte chemotaxis (GO:0048247, adjusted p< 0.005, includes *CCL19*, *CCL2*, *CCL13*), antigen presentation (GO:0019882, adjusted p< 0.005, includes *CD74*, *HLA-DRA*, *HLA-DRB1*), and positive regulation of T cell proliferation (GO:0042102, adjusted p< 0.005, includes *TNFSF13B*, *VCAM1*, *CCL5*). CXCL10^+^CCL19^+^ fibroblasts show broad evidence of response to key pro-inflammatory cytokines IFN_*γ*_ (GO:0034341, adjusted p=0.005), IFN*α* (GO:0035455, adjusted p=0.02), TNF*α* (GO:0034612, adjusted p< 0.005), IL-1 (GO:0070555, adjusted p< 0.005), and IL-12 (GO:0070671, adjusted p< 0.005). While TNF*α*, IL-1, and IL-12 response are broadly enriched in several fibroblast populations, an interferon response (IFN_*γ*_ and IFN_*α*_) is more specific to CXCL10^+^CCL19^+^ fibroblasts. In contrast to these cytokine-signaling pathways, SPARC^+^COL3A1^+^ fibroblast marker genes were enriched in pathways centered around extracellular matrix binding (GO:0050840, adjusted p< 0.005, includes *COL11A1, SPARC, LRRC15*) and disassembly (GO:0022617, adjusted p=0.005, includes *MMP13, MMP11, FAP*) and numerous developmental pathways (GO:0035904, GO:0060348, GO:0061448, GO:0007492, adjusted p< 0.005, includes *COL3A1, COL1A1, COL5A1, TGFB1*).

**Figure 5.**
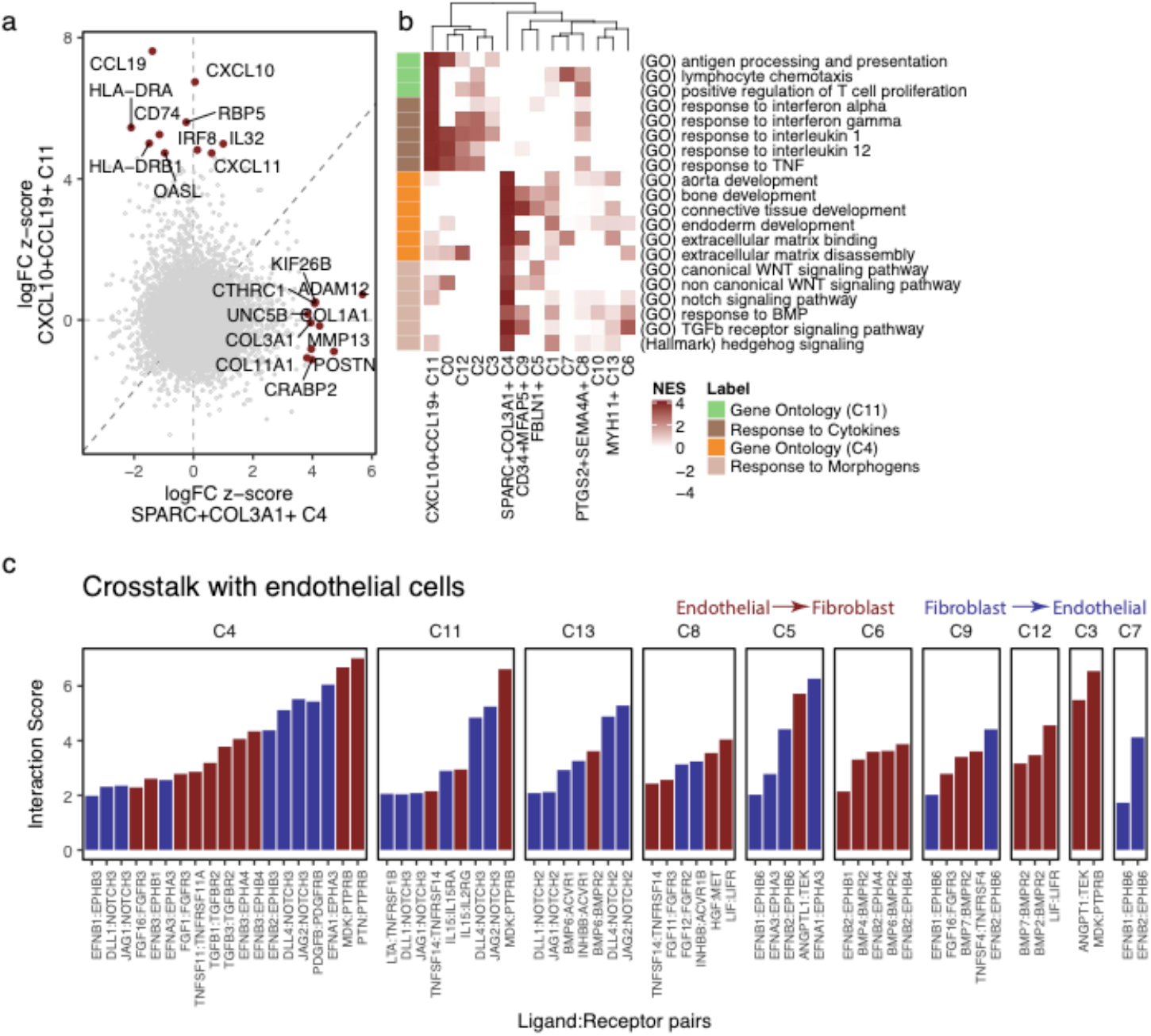
Distinct gene expression profiles for CXCL10^+^CCL19^+^ and SPARC^+^COL3A1^+^ states. (a) Comparison of differential gene expression between CXCL10^+^CCL19^+^ and SPARC^+^COL3A1^+^ fibroblasts shows that these two inflammation-expanded clusters are characterized by distinct genes. Top 10 markers for each cluster are named. (b) Gene set enrichment analysis with Gene Ontology and MSigDB Hallmark pathways shows distinct functions for the C4 (orange) and C11 (lime) states. These states may be explained by response to distinct sets of signaling molecules: inflammatory cytokines for C4 (brown) and tissue modeling morphogens for C11 (tan). Heatmap shows normalized enrichment scores from GSEA, focusing on only positive enrichment for clarity. (c) Ligand receptor analysis of endothelial cell crosstalk with fibroblast populations. Each column is a putative ligand receptor cognate pair, faceted by fibroblast subtype. Y-axis represents the strength of the putative crosstalk, while color denotes direction of interaction: (blue) endothelial ligand to fibroblast receptor or (red) fibroblast ligand to endothelial receptor.

Together, this suggests that SPARC^+^COL3A1^+^ fibroblasts may be driven by conserved developmental pathways during tissue remodeling in chronically inflamed diseases. Given the extensive enrichment in developmental pathways in these fibroblasts, we hypothesized that this state could be driven by morphogens within the tissue microenvironment. Indeed, we observed enrichment in key morphogen signaling pathways hedgehog (adjusted p=0.005), TGF*β* (GO:0007179, adjusted p< 0.005), WNT (canonical (GO:0060070, adjusted p=0.007) and non-canonical (GO:0035567, adjusted p=0.005)), BMP (GO:0071772, adjusted p=0.01), and Notch (GO:0007219, adjusted p< 0.005). Of these pathways, Notch signaling was the most specific to SPARC^+^COL3A1^+^ fibroblasts (**Figure 5b**), with non-significant (raw *p* > 0.20) enrichment in all other clusters. Since we have previously identified Notch3 signaling as a key driver in differentiation of disease-associated perivascular fibroblasts in RA synovia (Wei et al., 2020), we predict this cluster may represent a similar endothelium-driven, activated fibroblast state across inflammatory diseases involving other organ tissues. We explored this hypothesis with ligand receptor analysis (**Methods**). We started with manually curated cognate ligand and receptor pairs (Ramilowski et al., 2015) and for each pair, looked for high expression of one gene in endothelial cells within our libraries (**Figure 1b**) and its partner in each fibroblast state. Filtering for only differentially expressed genes, we found a total of 63 putative signaling interactions (**Figure 5c**). Notably, 19 of these interactions were between SPARC^+^COL3A1^+^ fibroblasts and endothelial cells, including Notch activation through the DLL4:NOTCH3 interaction, as described earlier in the synovium (Wei et al., 2020), as well as morphogen TGF*β*, growth factor PDGF*β*, angiogenic factors Ephrin-*α* and Ephrin-*β* (Rudno-Rudzińska et al., 2017), and angiogenic and mitogenic factors MDK and PTN (Weckbach et al., 2012). This large variety of putative signaling interactions (**Figure 5c**), both from and to endothelial cells, suggests that SPARC+COL3A1+ fibroblasts participate in signaling crosstalk with endothelial cells. Together, these pathway and crosstalk analyses suggest two independent, conserved populations that support tissue inflammation: namely immune cell-interacting CXCL10^+^CCL19^+^ immuno-fibroblasts and endothelium-interacting SPARC^+^COL3A1^+^ vascular associated fibroblasts.

### Correspondence between fibroblast clusters defined in integrative analysis and single-tissue analyses

We determined how the clusters labeled in the single-tissue analyses (**Figure 2a**) mapped to our new shared cross-tissue taxonomy. Since we used the same cells for both within-tissue and cross-tissue analyses, we were able to directly compare the overlap (**Methods**) between these two types of state definitions (**Supplementary Figure 5a**). The immuno-fibroblast cluster C11 overlapped significantly (*FDR* < 5%) with THY1^+^ sublining (*OR* = 3.8, 95% *CI*[2.2, 6.7]) and HLA-DRA^hi^ synovial fibroblasts (*OR* = 39.2, 95% *CI*[22.2, 69.0]), with CCL19^+^ fibroblasts in the salivary gland (*OR* = 9.1, 95% *CI*[6.3, 13.0]), with RSPO3^+^ (*OR* = 16.1, 95% *CI*[12.0, 21.7]) and WNT2B^+^Fos^hi^ (*OR* = 2.3 95% *CI*[1.7, 3.1]) fibroblasts in the intestine, and did not overlap significantly with any one cluster in the lung. Here, odds ratio refers to the probability of a cell being in a cross-tissue cluster (versus not), given that the cell belongs to some within-tissue clusters. The vascular-fibroblast cluster C4 was split between DKK3^+^ and THY1^+^ sublining fibroblasts in the synovium, mapped exclusively to myofibroblasts in the lung, split between inflammatory fibroblasts and myofibroblasts in the intestine, and mapped to CD34^+^ fibroblasts in the salivary gland. Notably, none of these associations was one-to-one. HLA-DRA^+^ synovial fibroblasts, CCL19^+^ salivary gland fibroblasts, and RSPO3+ and WNT2B^+^Fos^hi^ intestinal fibroblasts mapped to multiple clusters that were expanded in one or more tissues: C3 (lung and synovium), C2 (synovium), C12 (intestine), and C8 (salivary gland and synovium). Similarly, the myofibroblasts in the lung and intestine, as well as DKK3^+^ synovial fibroblasts mapped to both C13 and to vascular fibroblasts (C4).

Cluster C13 aligned strikingly with intestinal and pulmonary myofibroblasts. Although C13 contained cells from all tissues, it only expressed canonical myofibroblast genes *MYH11, MYL9, and ACTA2* in intestinal and pulmonary cells (**Supplementary Figure 5b**). While myofibroblasts are absent in synovium, synovial C13 cells may reflect an activated phenotype involved in tissue repair. This is supported by synovial specific upregulation of bone and cartilage reparative genes *TFF3, BMP6, HTRA1, and HBEGF* (**Supplementary Figure 5c**).

In the synovium and intestine, several clusters have previously been shown to be associated with distinct anatomical locations (Mizoguchi et al., 2018; Smillie et al., 2019; Zhang et al., 2019): PRG4^+^ synovial lining fibroblasts, THY1^+^ sublining synovial fibroblasts, WNT5B^+^ villus-associated fibroblasts, and WNT2B^+^ crypt-associated fibroblasts. Many of the integrated clusters we identified grouped along these anatomically defined lines. Clusters C0, C6, C10, and C12 were most associated with PRG4+ lining-associated synovial and WNT5B+ villus-associated gut fibroblasts, while clusters C1, C2, C3, and C8, mapped to THY1+ sublining-associated synovial and WNT2B+ crypt-associated gut fibroblasts. Except for cluster C8, these clusters that were strongly associated with anatomical locations in gut and synovium had fewer numbers of shared marker genes across tissues, potentially reflecting tissue-specific functions dictated by the specific anatomical constraints and physiological functions of the tissue.

FBLN1^+^ C5 and CD34^+^MFAP5^+^ C9 states mapped strongly to RSPO3^+^ intestinal, HAS1^+^PLIN2^+^ pulmonary and CD34^+^THY1^+^ synovial fibroblasts. The remaining cluster C7 did not map well to intestinal or synovial clusters. Subsequent analysis of marker genes within tissues suggested enrichment in doublets: epithelial markers *KRT7* and *ADGRF5* in lung and macrophage markers *C1QB, C1QA, and SPP1* in the salivary gland. This suggests that despite our best efforts to filter doublets during QC preprocessing, some contaminating doublets were retained. This makes further inference about cluster C7 less reliable.

### Validation in an alternative tissue: dermal fibroblasts in atopic dermatitis

As a proof of principle, we next explored whether the fibroblast states discovered in the four tissues could generalize to a tissue not explored in this study by examining cells from an independent dataset. We analyzed data from a study (He et al., 2020) of atopic dermatitis (AD), a chronic inflammatory condition of the skin (**Figure 6a**). The authors performed droplet-based scRNAseq on all cells from cryopreserved skin biopsies of 5 patients with AD (4 samples from skin lesions and 5 samples from skin outside of lesions) and 7 healthy donors. After removing low-quality (**Methods**) cells and 3 samples with fewer than 500 high-quality cells, we clustered 29,625 cells from 13 samples to identify the following major cell types (**Supplementary Figure 6a-b**): *MLANA*^+^ melanocytes, *KRT15*^+^ epithelial cells, *CD3G*^+^ T cells, *C1QB*^+^ myeloid cells, *PROX1*^+^ lymphatic endothelial cells, *ACKR1^+^* vascular endothelial cells, *ACTA2^+^* mural cells, and *COL1A1^+^* fibroblasts. As before, we used immune cell abundance to quantify a relative inflammation score in each sample (**Figure 6b**). Immune cell abundance correlated with histological classification, highest in samples from skin lesions and lowest in samples from non-diseased controls (**Figure 6b**).

**Figure 6.**
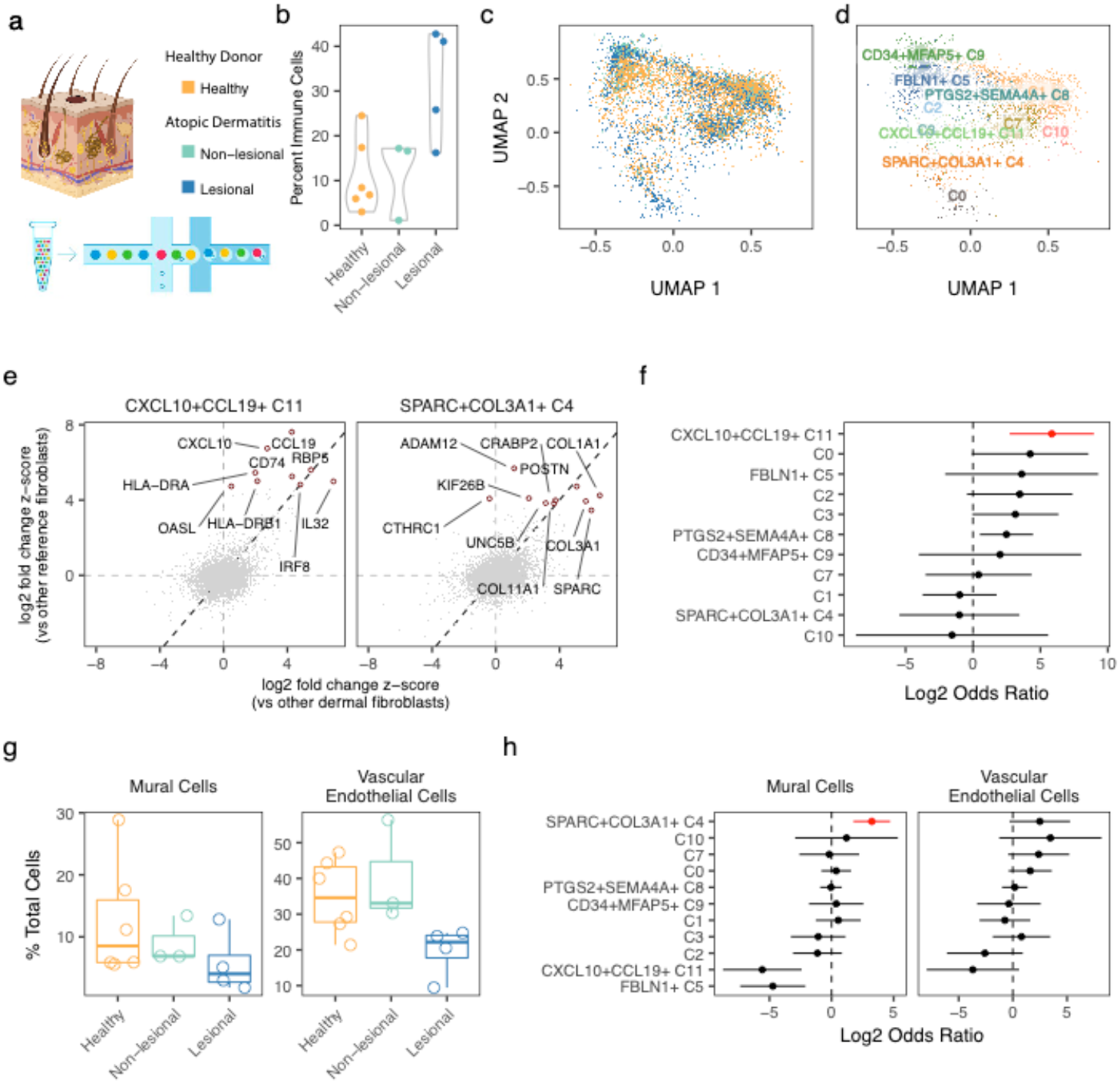
Dermal fibroblast scRNAseq profiles mapped to cross-tissue fibroblast atlas. (a) To validate our results, we mapped scRNAseq profiles of dermal fibroblasts from lesion biopsies from atopic dermatitis (AD) patients, non-lesional biopsies from AD patients, and control skin biopsies from healthy donors. (b) Based on the relative frequency of immune cells in each biopsy, we computed standardized inflammation scores from 0 to 1. (c) We mapped dermal fibroblasts to our fibroblast atlas and (d) labeled dermal fibroblasts according to their most similar atlas cluster. (e) We confirmed that the gene expression profiles of inferred dermal fibroblast clusters correlated with expression profiles of their reference fibroblast clusters. This is demonstrated for clusters C4 and C11 by plotting the (differential) gene expression in dermal (x-axis) vs reference (y-axis) clusters and calling out the top marker genes identified in the reference clusters. (f) Only CXCL10^+^CCL19^+^ (C11) fibroblast frequency was significantly (FDR<5%) associated with dermal inflammation. (g) Cells from skin with lesions (blue) had considerably less evidence of vasculature, measured by the abundance of perivascular mural cells and vascular endothelial cells. (h) Relative abundance of mural and endothelial cells was most strongly associated with cluster C4. Red denotes one-tailed FDR<5%.

We wanted to compare dermal fibroblasts directly to clusters defined in our fibroblast atlas. To do this, we leveraged a novel algorithm, Symphony (Kang et al., 2020) (**Methods**), designed to quickly and accurately map new scRNAseq profiles into a harmonized atlas to compare them with annotated reference cells. Using Symphony, we mapped dermal fibroblasts into our multi-tissue fibroblast atlas and projected them into the reference UMAP space for visual comparison (**Figure 6c**). For quantitative comparison of fibroblast subtypes, we labeled individual dermal fibroblasts by their most similar reference clusters (**Figure 6d**). Dermal fibroblasts mapped primarily to all clusters except C6, C12, and C13, three clusters which we identified as more tissue-specific (**Figure 3g**). We computed marker genes for these clusters in skin (**Supplementary Table 9**) and compared them to the markers we computed in the cross-tissue analysis. Encouragingly, the gene expression profile of each dermal fibroblast cluster most closely resembled that of its corresponding reference cluster (**Supplementary Figure 6c**). As two examples of this expression concordance, we plotted gene expression of immune (C4) and vascular (C11) fibroblasts inferred in the skin dataset versus those labeled in the reference (**Figure 6e**), highlighting the top 10 marker genes upregulated in each of the fibroblast clusters in the reference (**Figure 6e**).

We associated the abundance of inferred dermal fibroblast clusters with the sample-level inflammation score (**Figure 6f**). CXCL10^+^CCL19^+^ (C11) fibroblasts were the most significantly expanded in inflamed skin samples (*OR* = 57, 95% *CI* [6.5, 503], *p* = 2 × 10^−4^), even when performing the association within histological groups *OR* > 1000, *p* = 1.8 × 10^−11^) (**Supplementary Figure 6d**). Interestingly, SPARC^+^COL3A1^+^ fibroblasts, expanded in the original four tissues, were less abundant in inflamed skin. Given the previous association of SPARC^+^COL3A1^+^ fibroblasts with vasculature, we explored the relative degree of vascular cell types in each skin sample. Intriguingly, lesional samples had significantly fewer vascular endothelial (one-tailed t-test *p* = 0.004) and perivascular mural (one-tailed t-test *p* = 0.07) cells (**Figure 6g**), as compared to non-lesional and healthy samples together. The lack of vascular fibroblast expansion in inflamed samples from skin lesions is consistent with this decreased vascularization. In fact, the abundance of vascular fibroblasts is associated nominally with the abundance of vascular endothelial cells (log *OR* = 2.5, *p* = 0.04) and strongly with perivascular mural cells (log *OR* = 3.2, *p* = 1.8 × 10^−5^), when taking into account the histological status (**Figure 6h**).

### Cross-species mapping identifies shared fibroblast activation states in disease animal models of pulmonary, synovial, and intestinal inflammation

Next, we tested whether our two shared inflammation associated fibroblast subtypes were identifiable in single-cell datasets from mouse models of tissue inflammation. By defining which aspects of fibroblast-driven pathology are reproduced in mouse models, it may be possible to elucidate which pathological processes in murine models best parallel human fibroblast cell states. We found three single-cell RNAseq data sets that included both inflamed and non-inflamed samples in matched mouse tissues, which we could use to analyze both the conservation of cluster markers and the expansion of inflammation-associated immuno-fibroblasts and vascular fibroblasts (**Figure 7a**): Kinchen et al., 2018 profiled 8,113 cells, CD45^−^ gated to enrich for stroma, from 3 healthy and 3 mice with Dextran Sulfate Sodium (DSS)-induced colitis. Tsukui et al., 2020 profiled 15,095 cells, Col1a1^+^ gated to enrich for fibroblasts, from 2 healthy and 2 bleomycin-induced lung injury mouse lungs. Wei et al., 2020 profiled 8,738 total synovial cells from mice with K/BxN serum transfer induced arthritis, half with active inflammation and half with abated disease by inhibition of Notch3 signaling, by genetic knockout (Notch3^−/−^) and blocking antibody (anti-Notch3 mAB). Of note, while the K/BxN transgenic model generates autoreactive antibodies through a lymphocyte-mediated etiology, mice receiving those autoreactive antibodies through serum transfer develop arthritis through a lymphocyte-independent etiology (Monach et al., 2007). Therefore, we did not expect to see changes in the frequency of T cell interacting immuno-fibroblasts with this model.

**Figure 7.**
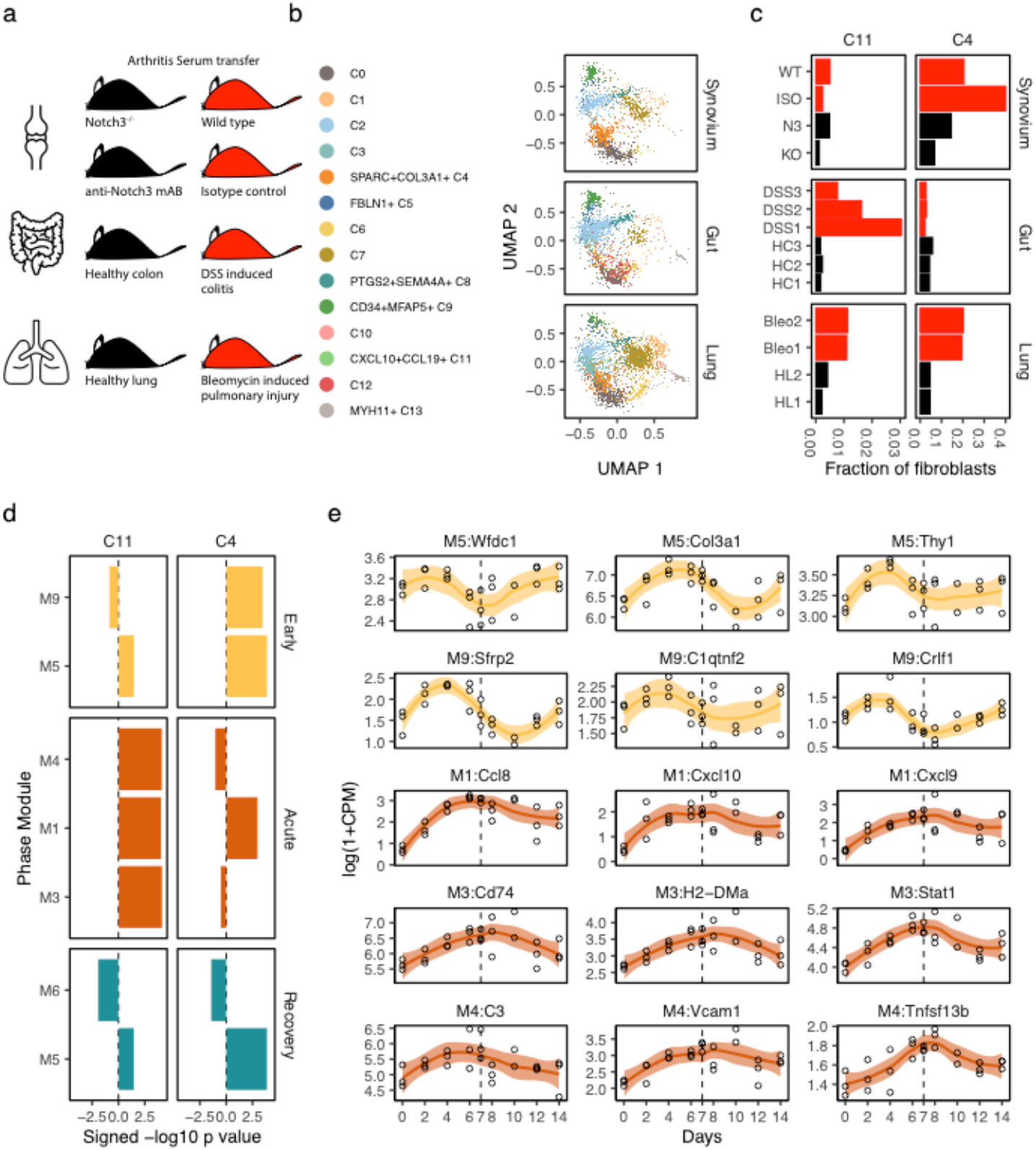
Replication in disease models of pulmonary, intestinal, and synovial inflammation. (a) We collected studies of inflammation in mouse models of human disease: bleomycin induced ILD, DSS-induced colitis, and serum transfer arthritis. (b) Fibroblasts from each study were mapped to the human fibroblast atlas and labeled with their most closely mapped clusters. (c) Frequencies of the human inflammatory states C4 and C11 in each study sample, colored to denote samples from animals with high (red) and low (black) inflammation. (d) Gene set enrichment analysis with modules associated with early, acute, and recovery phases of DSS-induced colitis shows that C4 and C11 gene signatures are activated at distinct stages of inflammation. (e) Time course expression profiles of key C4 and C11 marker genes that overlap with the early (yellow) and acute (orange) phase associated modules. Dotted line denotes timepoint (day 7) at which DSS was removed from mice.

Within each study, we identified fibroblasts (6,979 intestinal, 10,320 pulmonary, and 5,704 synovial) with clustering and marker analyses (**Supplementary Figure 7a,b**). We then mapped these fibroblasts to our human cross-tissue reference with the Symphony pipeline (**Methods**) and labeled mouse cells with the most similar reference fibroblast subtypes (**Figure 7b**). While most clusters were well-represented across tissues, two appeared more tissue-specific (**Supplementary Figure 7c**). Myofibroblast-enriched C13 was mostly absent in synovium, which is known to lack myofibroblasts. Cluster C12, which mapped well to the intestinal WNT5B^+^ 2 cluster in our initial analyses (**Supplementary Figure 5a**), was enriched in intestinal fibroblasts in this mouse analysis. To test the degree to which gene markers are conserved between mouse and human, we performed cluster marker analysis in the mouse fibroblasts (**Supplementary Table 10**) and compared cluster expression profiles between mouse genes and human orthologs (**Supplementary Figure 7d**). Importantly, the most similar gene expression profiles were between corresponding clusters in mouse and human. Moreover, for most clusters, expression profiles were even more similar between matched tissues.

We next asked whether the same fibroblast subtypes were expanded in inflamed tissues in human disease and mouse models. Thus, we performed differential abundance analysis within each mouse dataset (**Supplementary Figure 7e**), comparing inflamed cases to matched controls (**Methods**) to determine whether the SPARC^+^COL3A1^+^ and CXCL10^+^CCL19^+^ populations expanded in human tissues were also expanded in mouse models (**Figure 7c**). In bleomycin treated lungs, the most expanded populations were SPARC^+^COL3A1^+^ (*OR* = 5.2, 95% *CI* [4.5, 6.0], *p* < 10^−8^) and CXCL10^+^CCL19^+^ (*OR* = 3.8, 95% *CI*[2.2, 6.6], *p* = 2.5 × 10^−6^) fibroblasts. In arthritis models, the Notch signaling enriched (**Figure 5b**) SPARC^+^COL3A1^+^ cluster was greatly diminished with therapeutic Notch3 inhibition (*OR* = 3.8, 95% *CI*[1.5, 9.4], *p* = 4.1 × 10^−3^). On the other hand, the frequency of lymphocyte-interacting CXCL10^+^CCL19^+^ fibroblasts was not associated with disease activity in arthritic mice (*OR* = 1.2, 95% *CI*[0.47, 3.3], *p* = 0.6). This result is consistent with the known lymphocyte independence of the serum transfer model etiology (Monach et al., 2007). In DSS-induced colitis, CXCL10^+^CCL19^+^ fibroblasts were significantly expanded (*OR* = 6.1, 95% *CI*[1.9, 19.3], *p* = 2.3 × 10^−3^), as reported previously (Kinchen et al., 2018), while SPARC^+^COL3A1^+^ fibroblasts were actually diminished (*OR* = 0.5, 95% *CI*[0.4, 0.7], *p* = 9.2 × 10^−7^) in frequency.

### Temporal ordering of C4 and C11 activation in DSS-induced colitis

We were surprised that SPARC^+^COL3A1^+^ fibroblasts were not significantly expanded in a DSS-induced colitis model, despite their significance in the human cohorts. The lack of SPARC^+^COL3A1^+^ signal could mean that DSS-induced colitis utilizes an alternative inflammatory process. However, the difference may also reflect the kinetics of disease. Since DSS-induced inflammation is an acute process, reversible with removal of the chemical irritant, cross-sectional cellular compositions in that model may differ from compositions of chronically inflamed UC intestine. Specifically, if SPARC^+^COL3A1^+^ fibroblasts are responsible for tissue remodeling to enable leukocyte infiltration, then genes associated with SPARC^+^COL3A1^+^ fibroblasts should precede those associated with CXCL10^+^CCL19^+^ fibroblasts. To test this hypothesis, we used recently published time course transcriptional profiles of DSS-induced colitis, which tracks gene expression changes with the induction and resolution of inflammation (Czarnewski et al., 2019). The authors induced intestinal inflammation in female 8-12 week old C57BL/6J mice by putting DSS in their drinking water for 7 days and allowed resolution of inflammation by removing DSS for another 7. Measuring gene expression profiles with RNAseq approximately every 2 days, the authors defined gene modules M5 and M9 associated with early inflammation (2-4 days), M1, M3, and M4 with acute inflammation (6-8 days), and M5 and M6 with resolution (10-14 days). We analyzed the enrichment of these phase-associated modules in our fibroblast marker profiles to associate the expansion of fibroblast subtypes with distinct phases of DSS-induced inflammation and resolution (**Supplementary Table 11**). Strikingly, CXCL10^+^CCL19^+^ fibroblasts exclusively mapped to the three acute phase modules, M1, M3, and M4, while SPARC^+^COL3A1^+^ fibroblasts mapped to two early phase modules, M5 and M9 and only the M1 acute phase module (**Figure 7d**). Time course profiles of representative genes demonstrate the early and resolution phase activation of SPARC^+^COL3A1^+^-associated genes and acute phase activation of CXCL10^+^CCL19^+^-associated genes (**Figure 7e**). Given our hypothesis that SPARC^+^COL3A1^+^ fibroblasts are involved in vascular remodeling while CXCL10^+^CCL19^+^ fibroblasts interact with infiltrating immune cells, the early upregulation of SPARC^+^COL3A1^+^-association gene suggests that vascular remodeling precedes leukocyte infiltration in the DSS-colitis model.

## Discussion

In this study, we sought to define whether shared fibroblast states exist across four diverse tissues affected by clinically distinct inflammatory diseases. We postulated that defining shared pathogenic, inflammation-associated fibroblast states across diseases will help inform the possibility of common therapeutic strategies targeting fibroblasts across different inflammatory diseases. Comparison of pathogenic fibroblast phenotypes across diseases that manifest in different tissues is hampered by the lack of an accepted, tissue-independent taxonomy enjoyed by immune and vascular cells. We thus approached this question by generating novel single-cell RNAseq profiles of fibroblasts and analyzing the fibroblasts together to identify shared phenotypes across diseases. Cross-tissue analysis of gene expression is a challenging task, as evidenced by the plethora of statistical methods introduced to analyze even non-single-cell, multi-tissue data generated by the Genotype-Tissue Expression (GTEx) project (GTEx Consortium, 2015). By using sophisticated statistical methods for cross-tissue analysis, we were able to identify fibroblast phenotypes that were shared by all tissues as well as fibroblast adaptations unique to a subset of tissues.

The lack of universal definitions for key concepts such as fibroblast identity and inflammation scoring that apply equally well to all tissues presented a major challenge to our effort to associate fibroblast phenotypes with inflammation. In particular, the lack of a universal, pan-fibroblast surface marker prevented us from directly isolating fibroblasts with flow cytometry. We addressed this problem with negative selection, using specific markers to filter out non-fibroblast populations, and thus defining fibroblasts based on high-dimensional single-cell-RNA-seq data as non-epithelial, non-immune, non-endothelial, and non-mural cells with some known tissue-specific fibroblast markers, such as PDPN, PDGFRA, and COL1A1. The lack of a quantifiable score for inflammation impeded us from directly using standard tools from meta-analysis, which assume a standardized phenotype that can be measured equally well across all organ tissues. Inflammation in each disease is defined by disease-specific pathological processes, reflected in tissue-specific histological scores, such as the Krenn inflammation score in RA (Krenn et al., 2006) and Nancy index in UC (Marchal-Bressenot et al., 2017). We approached this challenge by intentionally selecting four chronic inflammatory diseases with distinct pathological and inflammatory processes. By analyzing fibroblasts from a range of diverse pathologies, we maximized the chances of identifying fibroblast phenotypes common to inflammation in four tissues. We chose the simplest aspect of inflammation that can be measured in all tissues, namely the proportion of immune cells infiltrating each tissue sample. Despite this simplicity, our definition robustly identified two shared fibroblast states, CXCL10+CCL19+ (C11) and SPARC+COL3A1+ (C4), associated with inflammation across tissues. A caveat of our definition of inflammation is that the other fibroblast clusters may be associated with distinct aspects of inflammation. For instance, PTGS2+SEM4A+ (C8) fibroblasts express neutrophil recruiting genes *CXCL1* and *CXCL2*, are critical to inflammation in UC (Friedrich et al., 2020), and likely associated with neutrophil infiltration.

The complexity of our study design, with cells measured from multiple donors, tissues, and diseases, presented a second major challenge to our study. Algorithms to identify shared clusters in scRNAseq datasets from multiple donors and tissues do not address key issues such as data imbalance or downstream analysis of gene expression in multi-tissue studies of human disease. Analyses that don’t account for these factors in this complex setting may result in diminished power and spurious associations. Here we use weighted PCA and weighted Harmony to account for imbalanced datasets and mixed effects Poisson regression to account for the effect of complex interactions between covariates on gene expression. Our analytical approach to decipher tissue-shared and tissue-specific gene expression serves as a template for well-powered and robust analysis of single-cell cluster markers, particularly relevant with the growing number of studies designed to identify shared etiology across tissues and diseases (Nieto et al., 2020; Szabo et al., 2019; Zhang et al., 2020).

Based on marker gene profiles, we believe that some of the clusters named in our analysis have been previously described in single-cell and functional studies of individual tissues, potentially with the exception of pSS, in which a scRNAseq atlas has not been described to date. For the first time, we provide a common frame of reference to cross-compare these diverse populations objectively across tissues. As a powerful corollary, we can draw upon functional studies performed in individual tissues to interpret the biological significance of our clusters.

CXCL10+CCL19+ (C11) fibroblasts closely resemble functionally well-characterized CCL19+PDPN+ immunofibroblasts in the salivary gland. These CCL19^+^ fibroblasts co-localize with CD3+ T cells and underlie the formation of salivary gland tertiary lymphoid structures in both human tissue and in an animal model (Nayar et al., 2019). This putative interaction with T cells is suggested by the expression of HLA genes in the synovial fibroblasts expanded in RA patients (Zhang et al., 2019). Here, HLA-DRA+ fibroblasts show strong evidence of response to IFN_*γ*_ and functional work demonstrated that IFN_*γ*_ is mostly produced by CD8+ T cells in inflamed synovium. Kinchen et al., 2018 also identified CCL19^+^ fibroblasts in the inflamed UC intestine, and numerous studies (Bisping et al., 2001; Breese et al., 1993) have associated T cells as the primary source of IFN_*γ*_ in intestinal inflammation. This suggests that T cell recruitment driven by CCL19+ fibroblasts and IFN-activated fibroblasts is a shared feature of inflammation across multiple diseases. Additional functional studies are required to investigate the complex interactions between T cells and fibroblasts in individual inflammatory diseases. Our integrative results provide generalizable markers that may identify such T cell interacting fibroblasts across tissues.

SPARC+COL3A1+ (C4) fibroblasts closely resemble the CD90^hi^ NOTCH3-activated synovial fibroblasts that are located near arterial blood vessels and pericytes and expanded in RA (Wei et al., 2020). Despite their perivascular location, NOTCH3^+^ fibroblasts, like our SPARC^+^COL3A1^+^ fibroblasts, are distinct from pericytes, as evidenced by their lack of canonical pericyte genes ACTA2 and MCAM (Armulik et al., 2011). Our cross-tissue analysis suggests that these vascular fibroblasts, which clustered separately from MCAM^+^ pericytes (**Figure 1**), may also play a role in vascular remodeling in the lung, intestine, and salivary gland. In the time-series analysis of acute inflammation in the mouse intestine, we found that the expansion of vascular fibroblasts preceded the expansion of CXCL10^+^CCL19^+^ immune-interacting fibroblasts. If this temporal ordering holds tissues, it suggests a two-stage mechanism for fibroblast-mediated regulation of inflammation, initiated by vascular remodeling that enables greater leukocyte infiltration into the tissue. Further mechanistic studies are needed to elucidate both the additional endothelium-derived, or angiocrine factors (Rafii et al., 2016) that mediate perivascular fibroblast differentiation and the mechanistic relationship between vascular and immune-interacting fibroblasts.

In interpreting clusters with more tissue-specific than tissue-shared genes, we noticed that tissue-specific programs often express genes with tissue repair functions. This observation may reflect the tissue-specific needs for maintenance and repair, defined by that tissue’s unique anatomical structures (Chang et al., 2002). In contrast, clusters with more tissue-shared genes were enriched in biological processes, such as immune cell recruitment (C11 and C8), processes which are independent of tissue architecture, and interaction with blood vessels (C4), structures which are present in all tissues. This dichotomy between functions tailored to a tissue’s structural composition versus functions common to all tissues explain why some fibroblasts phenotypes in scRNAseq appear more tissue-specific and others more tissue-shared.

We used a novel type of analysis from single-cell analysis called Symphony reference mapping (Kang et al., 2020) to compare human dermal fibroblasts and mouse lung, synovial, and lung fibroblasts to our annotated cross-tissue atlas. Reference mapping let us avoid intensive and error-prone manual interpretation steps in *de novo* analysis of the external datasets. We anticipate that this strategy can improve reproducibility in single-cell analysis in general and particularly in fibroblasts, whose phenotypes are often difficult to identify with one or two canonical marker genes. To promote reproducible research and cross-disease insights in fibroblast biology, we made both the fibroblast atlas (github.com/immunogenomics/fibroblastlas) and the tools needed to map data (github.com/immunogenomics/symphony) publicly available.

Fibroblasts are essential players in inflammatory disease, fibrotic disease, and cancer. The potential to target fibroblasts therapeutically is growing with the number of single-cell and functional studies on fibroblast heterogeneity (Dakin et al., 2018). While early studies of fibroblast heterogeneity focused on positional identity, more recent studies focus on functional states that mediate pathological processes. Our study provides the first cross-tissue analysis that rigorously distinguishes tissue-specific from tissue-shared identity in fibroblasts. In doing so, we described two fibroblast states that may be universal to inflammatory disease across tissues. In the process, we created the first single-cell reference atlas of fibroblast heterogeneity to unify fibroblast research and prevent a babelesque sprawl of fibroblast names across disciplines. Finally, we have proposed an analytical pipeline for studying shared pathological processes across diseases that can readily be applied to all cell types and tissues.

## Acknowledgements

We thank David Lee for having the vision and organizing this Roche network to study stromal biology across tissues. K.W. is supported by a NIH-NIAMS Clinical Investigator Award (1K08AR077037-01) and a BWH Department of Medicine Innovation Evergreen Award. B.A.F and S.J.B. have received support from the National Institute for Health Research (NIHR) Birmingham Biomedical Research Centre and the NIHR/Wellcome Trust Birmingham Clinical Research Facility. S.R. is supported by funding from the National Institutes of Health (U19AI111224, U01 HG009379, and R01AI049313). K.R. is supported by the NIHR Birmingham Biomedical Research Centre.

## Author contributions

I.K., K.W., and M.P. conceptualized study and co-wrote manuscript under supervision of S.R., M.B.B., C.D.B., and F.P. I.K. and J.B.K. performed analyses. K.W., M.P., E.Y.K., M.F., J.T., and S.N. performed experiments. E.Y.K., .A.F., K.R., F.B., B.A.F., S.J.B., C.D.B., and A.P.C. performed sample acquisition. All authors discussed results and commented on manuscript.

## Declaration of Interests

The authors have no declarations of interest to report.

## Data Statement

All FASTQ files and gene count matrices will be made available on NIAID ImmPort servers upon publication.

## Figure Legends

**Supplementary Figure 1. scRNAseq profiles of intestine, lung, salivary gland, and synovium.** (a) Flow sorting synovial and intestinal surgical samples to enrich for live (FVD^−^), EpCAM^−^CD45^−^ stromal cells. Cell level quality control summaries for scRNAseq libraries, represented with density plots of (b) percentage of mitochondrial reads and (c) the number of unique genes in a cell. (d) percentage of cells that were inferred to be doublets, of those that passed QC filtering (%MT≤ 20, nGene≥500). (e) Number of stromal and non-stromal cells identified in each tissue.

**Supplementary Figure 2. Labeling of previously defined fibroblast subtypes in each tissue.** Heatmaps represent the differential expression (one cluster vs all other clusters) z-scores of markers previously associated with published fibroblast subtypes in (a) synovium, (b) salivary gland, (c) intestine, and (d) lung. Columns (genes) colored by the fibroblast subtypes they are associated with.

**Supplementary Figure 3. Integrated cross-tissue fibroblast reference atlas.** (a) Breakdown of variance captured in the first 10 principle components for unweighted PCA and weighted PCA shows that weighted PCA creates a more balanced embeddings among tissues. (b) Before Harmony integration, UMAP embedding of fibroblasts separates entirely by tissue. (c) Within each tissue, there is substantial separation by donor, denoted by a different hue of the corresponding tissue’s color. UMAP coordinates are the same as in (b), zoomed in to focus on each tissue separately. (d) After Harmony integration, the clusters identified in tissue-specific analyses are still separated, suggesting that the Harmony embedding preserves within tissue variation. (e) Relative abundance integrative fibroblast clusters within each tissue.

**Supplementary Figure 4. Inflammation scores.** Comparison of differential abundance analysis using raw tissue-specific scores (x-axis) and normalized cross-tissue scores (y-axis). Error bars denote 95% confidence intervals.

**Supplementary Figure 5. Correspondence analysis.** (a) We associated cluster identity derived in single-tissue analyses (columns) to cluster identity derived in the integrative clustering analysis (rows). Color denotes (scaled) log odds from logistic regression. (b) Gene expression fold change of genes associated with myofibroblast lineage in cluster C13 (vs other clusters). (c) Same, for genes associated with bone and cartilage repair.

**Supplementary Figure 6. Dermal fibroblast scRNAseq profiles mapped to cross-tissue fibroblast atlas.** (a) UMAP embedding of scRNAseq profiles of skin biopsies, colored by major cell types, using (b) canonical markers: KRT15+ epithelial cells, COL1A1+ fibroblasts, PROX1+ lymphatic endothelial cells, MLANA+ melanocytes, C1QB+ myeloid cells, ACTA2+ mural cells, CD3G+ T cells, and ACKR1+ vascular endothelial cells. (c) Correlation of gene expression profiles of dermal fibroblast clusters (y-axis) against reference clusters in multi-tissue atlas (x-axis). Color denotes Pearson’s correlation coefficient. (d) Differential abundance of mapped dermal fibroblast clusters with inflammation score, with 95% confidence intervals. Red denotes FDR<5%.

**Supplementary Figure 7. Replication in disease models.** (a) UMAP embedding of mouse scRNAseq libraries from CD45^−^ sorted colon samples, unsorted synovial samples, and Col1a1^+^ sorted lung samples, colored by major cell types, identified with (b) canonical markers: Cdh5+ vascular endothelial cells, Col1a1+ fibroblasts, Lyve1+ lymphatic endothelial cells, Mcam+ mural cells, Myh11+ myofibroblasts, Ki67 proliferating cells, and Ptprc+ immune cells. (c) Relative abundance of inferred fibroblast clusters in each mouse dataset. (d) Comparison of mouse cluster gene expression profiles (y-axis) to human reference cluster profiles (x-axis). Heatmap color denotes Pearson’s correlation coefficient. Columns and rows are colored first by cluster identity and then by tissue. (e) Differential abundance of mapped mouse fibroblast clusters in case vs control mouse samples, with 95% confidence intervals. Red denotes FDR<5%.

**Supplementary Table 1.** Clinical characteristic for synovial tissue samples. Columns denote unique sample ID for each sample, clinical diagnosis, sex, age (in years), anatomical joint of surgical sample, and seropositivity status.

**Supplementary Table 2.** Clinical characteristic for lung tissue samples. Columns denote unique sample ID for each sample, clinical diagnosis, age (in years), sex, and serology.

**Supplementary Table 3.** Clinical characteristic for salivary gland tissue samples. Columns denote unique sample ID for each sample, sex, clinical diagnosis, presence or absence of anti-Ro antibodies, focus score, and free-text histology notes.

**Supplementary Table 4.** Clinical characteristic for intestine tissue samples. Columns denote unique sample ID for each sample, corresponding donor ID for repeat samples, histological status, year of birth, sex, Nancy score, and anatomical location of biopsy.

**Supplementary Table 5.** Cluster marker statistics for fibroblast cluster in single-tissue analyses. LogFoldChange is the differential expression of the gene (Feature) in the cluster (Cluster) against the mean of the remaining clusters within the tissue. Sigma is the estimated standard deviation around the log fold change statistic. Zscore is the standardized log fold change, divided by Sigma. Pval is the one tailed p value for the corresponding z score.

**Supplementary Table 6.** Association of inflammation score with pseudobulk fibroblast profiles. Columns same as in Supplementary Table 4, except for Slope, since inflammation score is a continuous and not a categorical covariate.

**Supplementary Table 7.** Cluster marker statistics for fibroblast subtypes defined in integrated analysis. Columns same as in Supplementary Table 4.

**Supplementary Table 8.** Gene set enrichment analysis of integrated fibroblast cluster markers. Columns are standard output of fgsea function. Pval is the nominal p value, padj is the adjusted p value, ES is the raw enrichment score, NES is the normalized enrichment score, nMoreExtreme is the number of more extreme observations in permutation tests, size is number of genes in the pathway, leadingEdge is the set of genes that contribute to the enrichment score.

**Supplementary Table 9.** Cluster marker statistics for dermal fibroblast subtypes. Columns same as in Supplementary Table 4.

**Supplementary Table 10.** Cluster marker statistics for mouse synovium, lung, and intestine fibroblast subtypes. Columns same as in Supplementary Table 4.

**Supplementary Table 11.** Gene set enrichment analysis of integrated fibroblast cluster markers. Columns same as in Supplementary Table 8.

## STAR Methods

### Human research and sample acquisition

Synovial study samples for transcriptomic studies were obtained from Brigham and Women’s Hospital, Hospital for Special Surgery, and the University of Birmingham under IRB-approved protocols. Synovial tissue from patients with clinically diagnosed rheumatoid arthritis were obtained from ultrasound-guided joint biopsy (University of Birmingham) or arthroplasty or synovectomy procedures (Brigham and Women’s Hospital and Hospital for Special Surgery). For arthroplasty and synovectomy tissue samples, the diagnosis of rheumatoid arthritis was confirmed clinically through clinical chart review. Synovial tissue from patients with osteoarthritis were obtained from arthroplasty procedures. Synovial tissues were cryopreserved on-site in Cryostor CS10, then shipped to BWH under a BWH IRB-approved protocol PROSET for tissue dissociation and single-cell transcriptomic analysis.

Intestinal samples were obtained from Ulcerative colitis (UC) or from healthy individuals by endoscopic biopsy. Healthy patients were recruited as a part of the research tissue bank ethics 16/YH/0247 and Inflammatory Bowel Diseases (IBD) patients among the Inflammatory Bowel Cohort 09/H1204/30 by the Translational Gastroenterology Unit Biobank at the John Radcliffe Hospital in Oxford. All patients gave informed consent and collection was approved by NHS National Research Ethics Service. Samples were immediately placed on ice (RPMI1640 medium) and processed within 3 hours.

Labial minor salivary gland samples were obtained from patients recruited in the Optimising Assessment in Sjögren’s Syndrome (OASIS) cohort (Machowicz et al., 2020) which recruits new patients attending the multidisciplinary Sjögren’s clinic at the Queen Elizabeth Hospital Birmingham, UK for assessment. Sjögren’s syndrome patients had a physician diagnosis of primary Sjögren’s syndrome and fulfilled the 2016 ACR/EULAR classification criteria. Participants with non-Sjögren’s sicca syndrome had signs and/or symptoms of dryness but did not have a physician diagnosis of SS or fulfill 2016 classification criteria. Salivary gland biopsy samples were divided in two: one for the scRNAseq study and the second for histological analysis to confirm diagnosis. Histological diagnosis is summarized in **Supplementary Table 3** and reported as presence of focal lymphocytic sialadenitis (FLS, suggestive of Primary Sjögren’s Syndrome, PSS) or non-specific chronic sialadenitis (NSCS), in the case of non-Sjögren’s sicca syndrome. Focus score (FSC, number of inflammatory foci/4mm^2^ of tissue) is also reported in Table 1. All OASIS participants provided written informed consent and the study was approved by the Wales Research Ethics Committee 7 (WREC 7) formerly Dyfed Powys REC; 13/WA/0392.

Lung samples were obtained from patients recruited at the Brigham and Women’s Hospital with informed consent under protocols approved by the Mass General Brigham IRB (PROSET). As enumerated in Supplementary Table 2, samples coded Lung1-15 (control donor lung, IPF, Rheumatoid Arthritis [RA]-ILD) were explants from lung transplant surgery. Samples coded Lung 16-23 (unclassifiable (u)ILD, IPF, NSIP) were from Video-assisted thoracoscopic surgical (VATS) lung biopsies for diagnosis of ILD. The patient condition is the diagnosis determined by clinical providers after their inter-disciplinary review of patient history, exam, clinical laboratory testing (e.g., serologies), imaging and histopathology of the explanted or biopsied lung tissue. The presence or absence of anti-CCP antibodies is noted.

### Cell isolation for single-cell RNA-sequencing

Synovial tissues were cryopreserved on site, thawed and disaggregated into single-cell suspension as previously described (Donlin et al., 2018). Four pairs of intestinal biopsies were pooled, minced and frozen in 1mL of CryoStor® CS10 (StemCell Technologies) at −80°C then transferred in LN2 within 24 hours. Single-cell suspensions from these endoscopic biopsies were then prepared by thawing, washing and subsequent mincing of the tissue using surgical scissors. Minced tissue was then subjected to rounds of digestion in RPM-1640 medium (Sigma) containing 5% Fetal Bovine Serum (FBS, Life Technologies), 5mM HEPES (Sigma), antibiotics as above, and Liberase TL (Sigma), with DNAse I. After 30 minutes, digestion supernatant was taken off, filtered through a cell strainer, spun down, and resuspended in 10ml of PBS containing 5% BSA and 5mM EDTA. Remaining tissue was then topped up with fresh digestion medium until no more cells were liberated from the tissue. Cells were then stained and FACS-sorted for live EPCAM^−^CD45^−^ cells, before being taken for microfluidic partitioning.

Lung tissues were cryopreserved on site, thawed and disaggregated into single-cell suspension. Each lung tissue was frozen in 1mL of CryoStor CS10 in −80°C with a controlled rate of freezing and then transferred to LN2 within two weeks. On the day of single-cell analysis, the cryopreserved lung tissue was rapidly thawed, serially rinsed with DMEM (GIBCO) supplemented with 10% FBS and then DMEM with 2% FBS on ice. Lung tissue was minced using surgical scissors and then transferred to a polypropylene tube with digestion media containing Liberase TL, hyaluronidase (Worthington Biochemical Corporation), Elastase (Worthington Biochemical Corporation), DNAse (Sigma) and 1% FBS. The addition of FBS improved cell viability without reducing yield of viable stromal cells. After 20 minutes of incubation at 37°C warm room with agitation by stir bar, the supernatant containing single cells was collected, and fresh digestion media was added. After 20 minutes of addition digestion, the tissue and supernatant were filtered through a 70 micron cell strainer and washed in DMEM with 2% FBS twice. Dead cells were removed using a magnetic column based method per manufacturers protocols (Dead Cell Removal kit, Miltenyi Biotec). Then single cells were taken for microfluidic partitioning.

Minor salivary gland biopsies were taken surgically from the lip and frozen in 1mL of CryoStor® CS10 (StemCell Technologies) at −80°C. For preparation of single-cell suspension, firstly the frozen tissue sample in Cryotube were quickly thawed in water bath at 37°C and washed twice in pre-warmed 5%FBS RPMI media. The salivary gland biopsies were then enzymatically digested as previously described (PMID: 31213547). Dead cells were removed using the EasySepTM Dead Cell Removal (Annexin V) kit from the digested samples following manufacturer’s instructions before proceeding for the scRNA sequencing using the 10x platform.

### RNA-sequencing

Single-cell RNA-sequencing experiments for lung, intestine, and synovium samples were performed through the Brigham and Women’s Hospital Single Cell Genomics Core. Viable cells in single-cell suspension were resuspended in 0.4% BSA in PBS at a concentration of 1,000 cells per ul. 7,000 cells were loaded onto a single lane (Chromium chip, 10X Genomics) followed by encapsulation in lipid droplet, with the 10x Genomics Single-Cell 3’ kit (Version 2 for synovium and intestine, Version 3 for lung) followed by cDNA and library generation per manufacturer protocol. cDNA libraries were sequenced to an average of 50,000 reads per cell using Illumina Nextseq 500. Single-cell RNA-sequencing experiments for salivary gland samples were performed at Oxford University. For each library, 10,000 cells were counted using the automated cell counter Bio-Rad TC20 and loaded onto a single 10x lane and processed with the 10x Genomics Single Cell 3’ kit (Version 3). Sequencing was done using Illumina NovaSeq 6000 and libraries were sequenced to a minimum of 50000 reads/cell.

### scRNAseq gene quantification

For all scRNAseq datasets analyzed in this manuscript, we quantified gene expression *ab initio* from FASTQ files. Human reads were mapped to the GRCh38 (Schneider et al., 2017) reference and genes annotated with Gencode (Frankish et al., 2019) v33. Mouse reads were mapped to mm10 reference and genes annotated with Gencode v25. For both human and mouse data, we filtered transcripts for the annotation “protein_coding” and ignored the rest. Reads from distinct transcripts of the same gene were collapsed by summation. We used kallisto (Bray et al., 2016) v0.46.0 to map reads to transcriptomes and bustools (Melsted et al., 2019) v0.39.3 to collapse duplicate reads by UMI and return gene-cell count matrices. We downloaded read level data for the following publicly available scRNAseq datasets: PRJNA614539 (He et al., 2020) (atopic dermatitis), PRJNA542350 (Kinchen et al., 2018) (DSS model), and PRJNA548947 (Tsukui et al., 2020) (Bleomycin model). After contacting the authors, the PRJNA542350 data turned out to be BAM files rather than FASTQ. Per their suggestion, we used the 10X Cell Ranger (Zheng et al., 2017) bamtofastq utility (version 1.3.2), with default parameters, to convert the BAMs back into FASTQs for remapping. doc. The code to perform all steps of this mapping are implemented as functions in the github repository for this manuscript.

### scRNAseq quality control, pre-processing, and normalization

After quantifying gene count matrices with kallisto and bustools (above), we filtered out poor quality cells with three metrics. (1) Cells must have at least 500 unique genes. (2) Cells must have more than 20% of the total UMIs mapped to non-mitochondrial genes. (3) Cells must be inferred as singlets by algorithmic doublet identification. For doublet identification, we used the scDblFinder algorithm(Germain, 2020), with default parameters, separately within each 10X library. We normalized for read depth with the standard logCP10K normalization procedure for gene *g* and cell 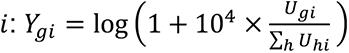.

### Inflammation score normalization across tissues

Inflammation scores computed within each tissue had ranges and distributions. To be able to compare inflammation associated phenotypes across tissues, we normalized the distributions by performing quantile normalization. Because the number of samples was relatively small, we did not use an empirical distribution. Instead, we normalized to the quantiles of a parametric distribution. We chose the beta distribution (*α* = 3, *β* = 3) to map the scores to an interpretable interval, between 0 (low inflammation) and 1 (high inflammation).

### Gene selection

For analyses with one tissue, we used the VST method for variable gene selection, reimplemented from the Seurat package (Butler et al., 2018) as a stand along function in our github at immunogenomics/singlecellmethods. We used default parameters and kept the top 2000 genes, ranked by standardized variance. For the multi-tissue integrated analysis, we used genes that we found informative in at least one of the tissue-specific analyses of lung, salivary gland, intestine, and synovium. We defined informative genes with two analyses. The first analysis is differential expression of cluster-markers for tissue-specific fibroblast subtypes (**Figure 4a**). We kept cluster-informative genes with *p* < 0.05 and |*β*| ≥ 0.5. The second analysis found broadly inflammation associated genes by fitting a Poisson log-normal GLMMs to each gene. We kept inflammation associated genes with *p* < 0.05 and |*β*| ≥ 0.1.

### Weighted PCA

We implemented principle components analysis that gives equal weight to each tissue while preserving the total cell number (∑*_i_ w_i_* = *N*). The weights given to each cell were determined to meet this equal weight condition. These weights were then used in the scaling and SVD steps. For scaling, we computed weighted means and variance with the following formulas: 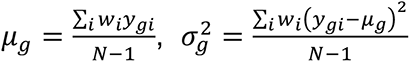. For SVD, we modified the PCA covariance decomposition formula to allow for observation weights with a diagonal matrix *W*: *XWX^T^* = *UDU*^*T*^. This decomposition is achieved by performing SVD on the weighted matrix *XW*^1/2^ *UDV*^*T*^. Because *W* is diagonal, its square root is the element-wise square root. This SVD solution now represents the original data as *X* = *UDV*^*T*^*W*^−1/2^, with gene loadings *U* and cell embeddings. Weighted PCA is implemented on our github at immunogenomics/singlecellmethods with the weighted_pca function.

### Weighted Harmony

We modified the Harmony algorithm to include observation weights. To achieve this, we modified the clustering objective function and rederiving the optimization steps for this function. The new objective function modifies the original only by multiply the per-cell cost (inside the summation) by 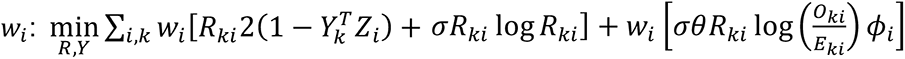. The rest of the formula is unchanged and described in detail in the original Harmony manuscript (Korsunsky et al., 2019). This modified Harmony implementation is available on our github at immunogenomics/harmony, under the weights branch.

### UMAP visualization

We used the UMAP algorithm to visualize cells in two dimensional embeddings. We used the uwot R package (Melville, 2020) with parameters n_neighbors=30L, metric=’Euclidean’, init=’Laplacian’, spread=0.3, min_dist=0.05, set_op_mix_ratio=1.0, local_connectivity=1L, repulsion_strength=1, and negative_sample_rate=1. For all other parameters, we used default values. In the symphony pipeline, we visualized mapped query cells by using the UMAP object learned for the reference analysis. The umap reference projection was done with the umap_transform function in uwot.

### Clustering

We performed graph based clustering with the Louvain algorithm (Blondel et al., 2008), implemented in Seurat (Butler et al., 2018). Instead of constructing the kNN and sNN graphs from scratch, we used the uniform manifold graph estimated in the UMAP algorithm. In the uwot package(Melville, 2020), this data structure is directly available in the fgraph field when umap is run with option ret_extra = c(‘fgraph’).

### Hierarchical gene expression modeling

#### Statistical model

We modeled the expression of each gene using Poisson lognormal GLMM regression. This framework allows us to model the hierarchical design in our multi-tissue, multi-donor dataset. We fit the following GLMM for the integrated, multi-tissue analysis, regressing to the frequency of gene *g* in observation *i*.

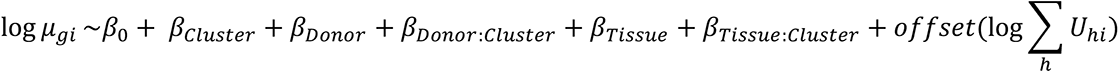

We chose to model the cluster interaction terms with donor and tissue. As many papers have observed (Haghverdi et al., 2018; Korsunsky et al., 2019), the effect of biological and technical covariates are often cell type specific. This is why integration algorithms cannot adjust every cell type by the same amount to account for batch, donor, or tissue variability. Unfortunately, the absence of some donors and tissues in some clusters means that interaction terms may be very poorly estimated. To address this issue, we model all terms except for the global intercept (*β*_0_) with Gaussian priors, allowing each effect to have a different size, denoted by τ^2^, the variance of the priors. These priors shrink *β*s towards zero, stabilizing estimation for terms with little data to draw from.

We performed cluster marker analysis with the estimated *β*s, estimating both marginal effects and tissue-specific effects. *Marginal cluster effects* are only concerned with the *β_cluster_* term. For instance, the differential expression for cluster 3 is 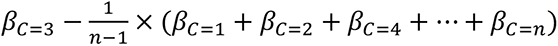. This comparison can be compactly represented with the contrast vector 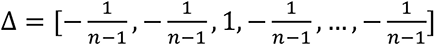 such that the differential expression can be computed with the linear operation 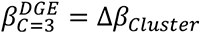. Following the example of significance testing in DESeq2, the standard errors of contrasts are in the diagonal elements of 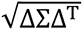, in which Σ is the covariance matrix of *β* levels. In our example, Σ is a cluster by cluster covariance matrix and the standard error for cluster 3 would be 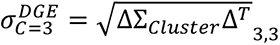. There is generally no analytical way to compute Σ for random effects, so we estimate it with simulation, using the arm R package(Gelman and Su, 2020), with 1000 simulations. *Tissue-specific cluster effects* take into account both the cluster and tissue-cluster interaction term. For instance, if we wanted to know how a gene is associated with cluster 3 in the lung, we would compute 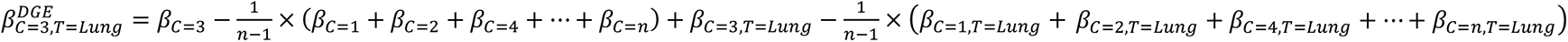. The contrast vector now includes terms that represent the *β*s estimated for lung tissue as well. The statistical procedures to compute 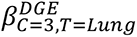 and 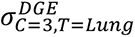 are the same as before. For both marginal and tissue-specific effects, we use a Gaussian approximation to estimate p values for each effect: 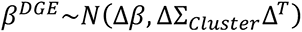.

#### Implementation

We fit GLMMs with the glmer function in the lme4 R package (Bates et al., 2015) and estimated random effect covariance with the sim function in the R arm package(Gelman and Su, 2020). Initially, we found it difficult to tie model fitting and simulation seamlessly with differential expression analysis. For instance, building contrasts for nested effects and estimating significance for multiple gene queries was difficult to do. Moreover, the memory footprint of lme4 models makes it impractical to fit and save models for 1000s of genes for downstream inference. To make lme4 and arm more accessible for gene expression analysis, we created the Presto package. Presto extracts the necessary components from lme4 models, saves them in efficient data structures, and has all necessary functions to do efficient contrast analysis for differential expression. We made Presto available as an R package, available on github at immunogenomics/presto under the GLMM branch.

To make the models more numerically stable, we enforced a minimum value for the size of random effects: σ ≥ 0.5. This prevented degenerate solutions with σ = 0, local minima which may arise in GLMM optimization. As a side effect, this Bayesian variance prior also enforces a conservative null model on random effects, effectively setting the null effect size to 0.5 rather than 0. This results in higher estimated uncertainty thus more conservative p values. In developing this software, QQ plot analysis was deflated and resembled post-hoc adjusted (e.g. Bonferroni) p values more than nominal p values from independent tests. Others have noted a similarity between *post hoc* correction and shrinkage integrated into the model (Gelman et al., 2012). For our analyses, we consider significance with respect to these shrunken p values, estimated with random effects, without doing additional *post hoc* shrinkage.

We made two decisions to make Presto scale to large datasets. First, we fit the model with pseudobulk, rather than single-cell RNAseq profiles. Note that in the formula above, the cluster, tissue, and donor covariates are not unique to single cells. Therefore, we collapse reads from cells with same cluster, donor, and tissue identity into one observation. This approach has strong precedent(Lun and Marioni, 2017). It is important to note that in this strategy, the number of parameters to estimate is equal to the number of observations. With fixed effects, this model is under-determined. However, because we shrink estimates to 0 with Gaussian priors, the effective number of independent parameters shrinks too. The second decision is with the choice of generative model. Many RNAseq differential expression tools used the Negative Binomial distribution, which uses Gamma rather than lognormal priors to model over-dispersion. For completeness, we also included negative binomial GLMMs in Presto. In practice, we found that this error model yielded almost identical results but took ten times longer to run.

#### Tissue heterogeneity

We took a very simple approach to labeling genes as conserved or heterogeneous cluster makers. Conserved markers were significantly (*p* < 0.05) overexpressed (*β* > 0) in all four tissues. If a gene was not upregulated in at least one tissue, we considered it to be a heterogeneous marker. Effect heterogeneity has a rich statistical treatment, especially in meta-analysis. We decided to not use these more sophisticated techniques, although the parameters learned in Presto could be used for such analyses.

#### Analyses

To find marker genes for dermal fibroblasts, we fit the same model as above but omitted the Tissue terms: 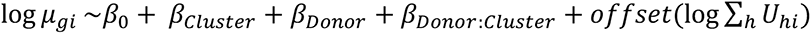. For the mouse scRNAseq analyses, we used the same hierarchical formula with all Tissue terms.

### Pathway analysis

All formal geneset enrichment was done with the GSEA algorithm, implemented in the fgsea R package(Sergushichev, 2016). To enrich pathways for marker analyses (**Figure 5d**), we used the H (hallmarks) and C5 (Gene Ontology) genesets from MSigDB, accessed with the msigdbr R package(Dolgalev, 2018). To enrich for different phases of inflammatory response in DSS-induced colitis (**Figure 7e**), we used the published genesets, provided as supplemental materials in the manuscript (Czarnewski et al., 2019).

### Abundance modeling

We associated inflammation score with cluster abundance using logistic regression, following the MASC method (Fonseka et al., 2018), with the following formula: 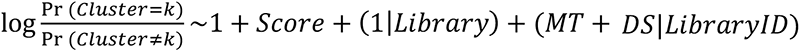. As in MASC, the response variable models the log odds of being in cluster *k* vs not, to test for which factors contribute to cluster *k* abundance. This probability is a function of (1) an intercept, which reflects the average abundance of cluster *k* in the data, (2) fixed effect for *Score*, the normalized inflammation score for each sample, (3) random effect for 10X library, to account for dependence of cells within a library, and (4) cell quality statistics *MT* (percent mitochondrial reads) and *DS* (doublet score), separately within each library. The association between inflammation and cluster abundance is captured in the *β* statistic. We computed significance for each *β* with the following Gaussian approximation, using the standard error σ provided by lme4: *β*∼*N*(0, σ^2^). To combine MASC results from individual tissue analyses, we used inverse variance weighted meta analysis with random effects. The variance from random effects was estimated with the DerSimonian and Laird (DL) method (DerSimonian and Laird, 1986; Veroniki et al., 2016).

### Cluster correspondence analysis

To compare the co-occurrence of the fibroblast cluster labels, within-tissue (**Figure 3**) and integrative (**Figure 4**), we used a similar framework to abundance modeling above. We used the following formula: 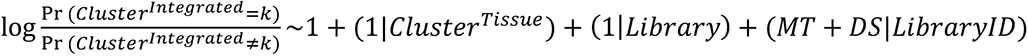. The contrast term of interest is the random effect (1|*Cluster^Tissue^*), a categorical variables that encodes the within-tissue cluster identity. We chose to model this with a random effect for numerical stability. To estimate significance, we used Wald’s approximation and simulated covariance for the levels of (1|*Cluster^Tissue^*) with the R arm package.

### Symphony projection

The Symphony pipeline is described in detail in a separate manuscript (Kang et al., 2020). In order to infer reference cluster identity in query cells, we used a k-NN classifier. K=10 nearest neighbors were estimated with Symphony projected low dimensional embeddings, based on cosine distance (σ = 0.1).

### Ligand receptor analysis

We started with a curated list of known interacting ligand-receptor pairs, from Ramilowski et al., 2015. To predict putative interactions between endothelial cells and fibroblast subsets, we performed differential expression on the pooled dataset of endothelial cells and fibroblasts. We filtered for differentially expressed genes and kept interaction pairs in which the ligand was overexpressed (*p* < 0.05, *β* > 0) in endothelial cells and the receptor in a fibroblast subset, or vice versa. For these pairs, we computed the interaction scores (**Figure 4e**) as the mean of the ligand’s and receptor’s z-scores.

## Notes

### Competing Interest Statement

The authors have declared no competing interest.

### Summary of Updates

Fixed supplementary table 7.

