## SupplementaryFigures for "Cross-tissue, single-cell stromal atlas identifies shared pathological fibroblast phenotypes in four chronic inflammatory diseases"

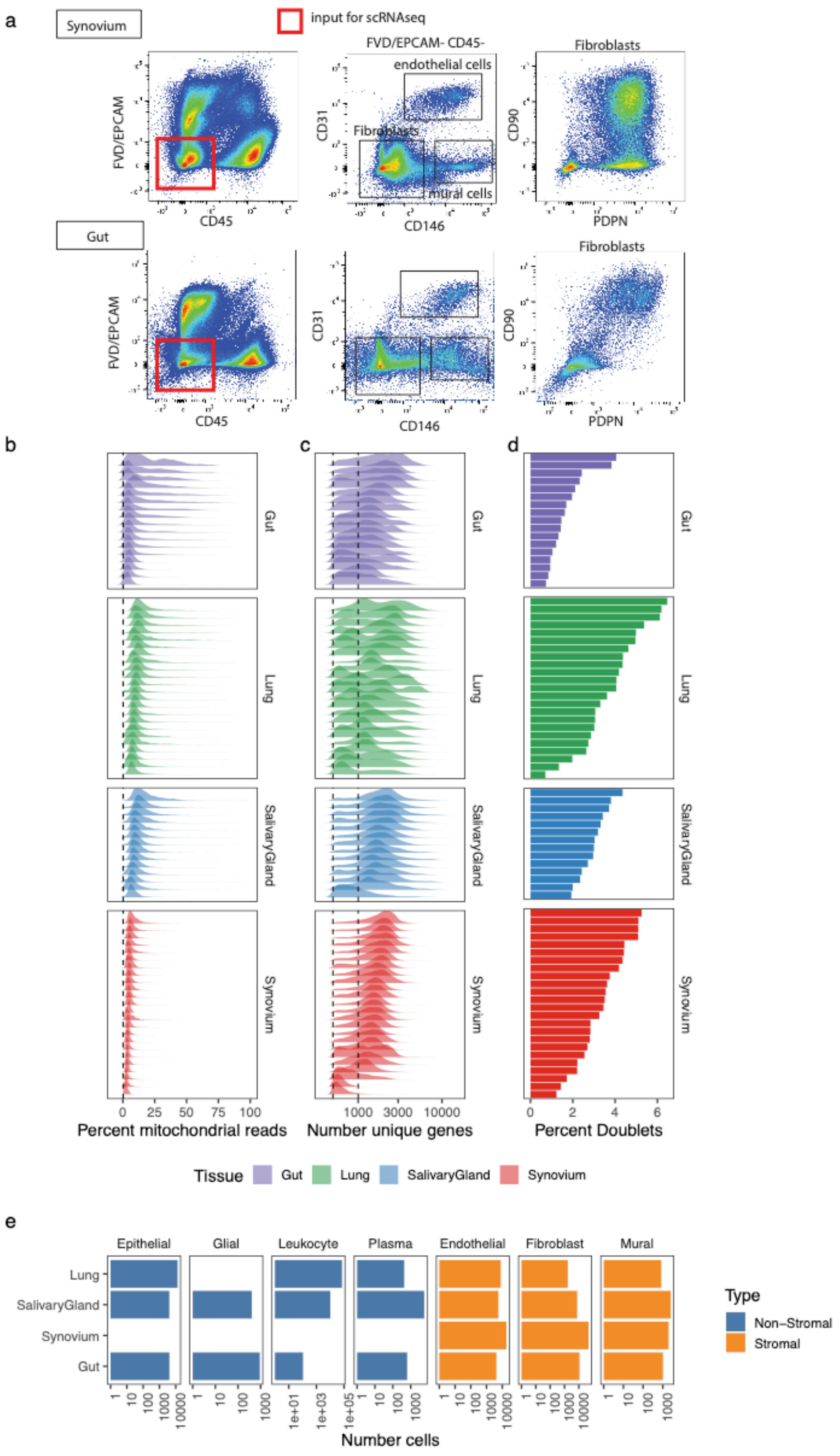


**Supplementary Figure 1. scRNAseq profiles of intestine, lung, salivary gland, and synovium.** (a) Flow sorting synovial and intestinal surgical samples to enrich for live (FVD$^-$), EpCAM$^-$CD45$^-$ stromal cells. Cell level quality control summaries for scRNAseq libraries, represented with density plots of (b) percentage of mitochondrial reads and (c) the number of unique genes in a cell. (d) percentage of cells that were inferred to be doublets, of those that passed QC filtering (%MT≤ 20, nGene≥500). (e) Number of stromal and non-stromal cells identified in each tissue.

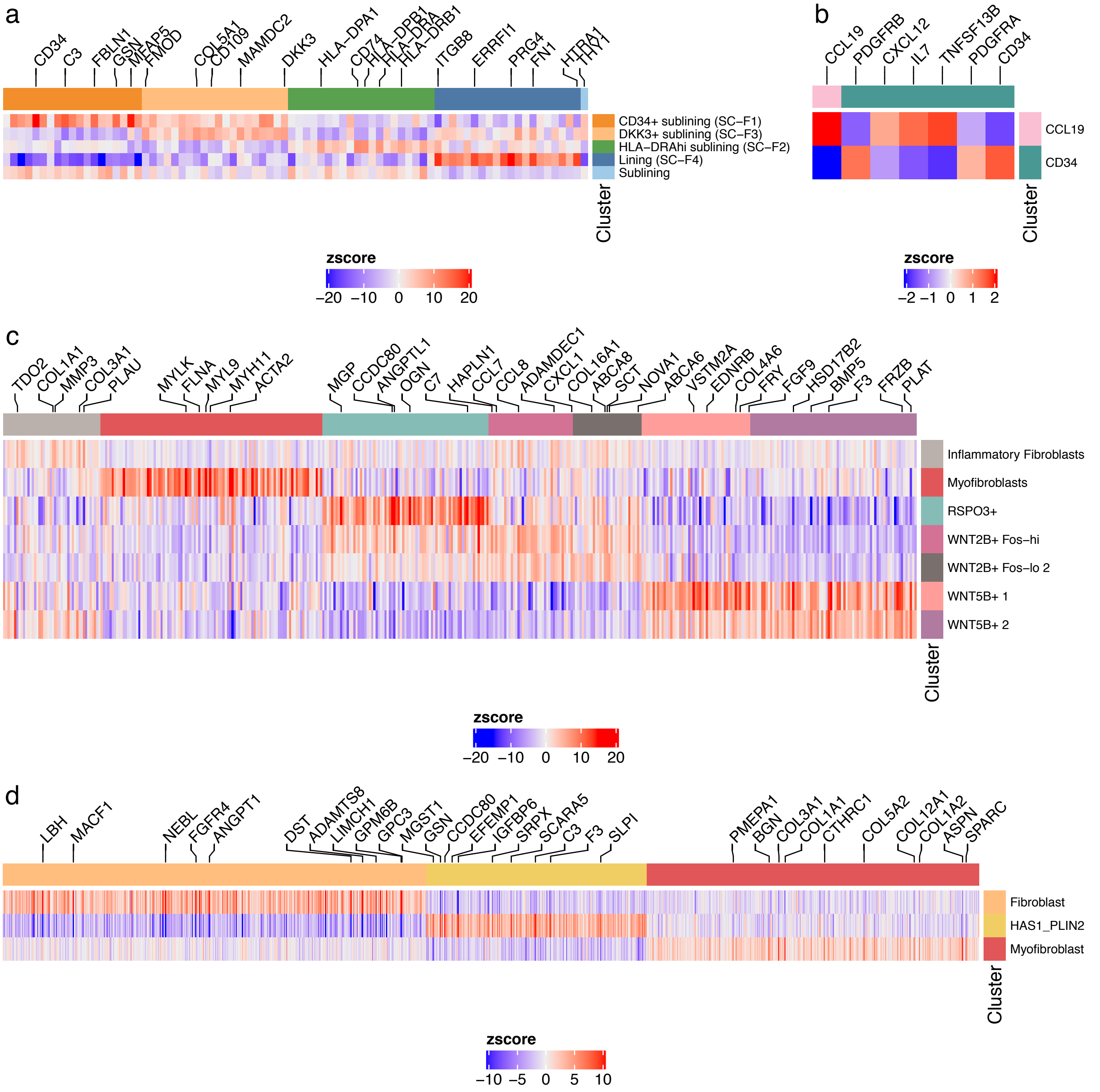


**Supplementary Figure 2. Labeling of previously defined fibroblast subtypes in each tissue.** Heatmaps represent the differential expression (one cluster vs all other clusters) z-scores of markers previously associated with published fibroblast subtypes in (a) synovium, (b) salivary gland, (c) intestine, and (d) lung. Columns (genes) colored by the fibroblast subtypes they are associated with.

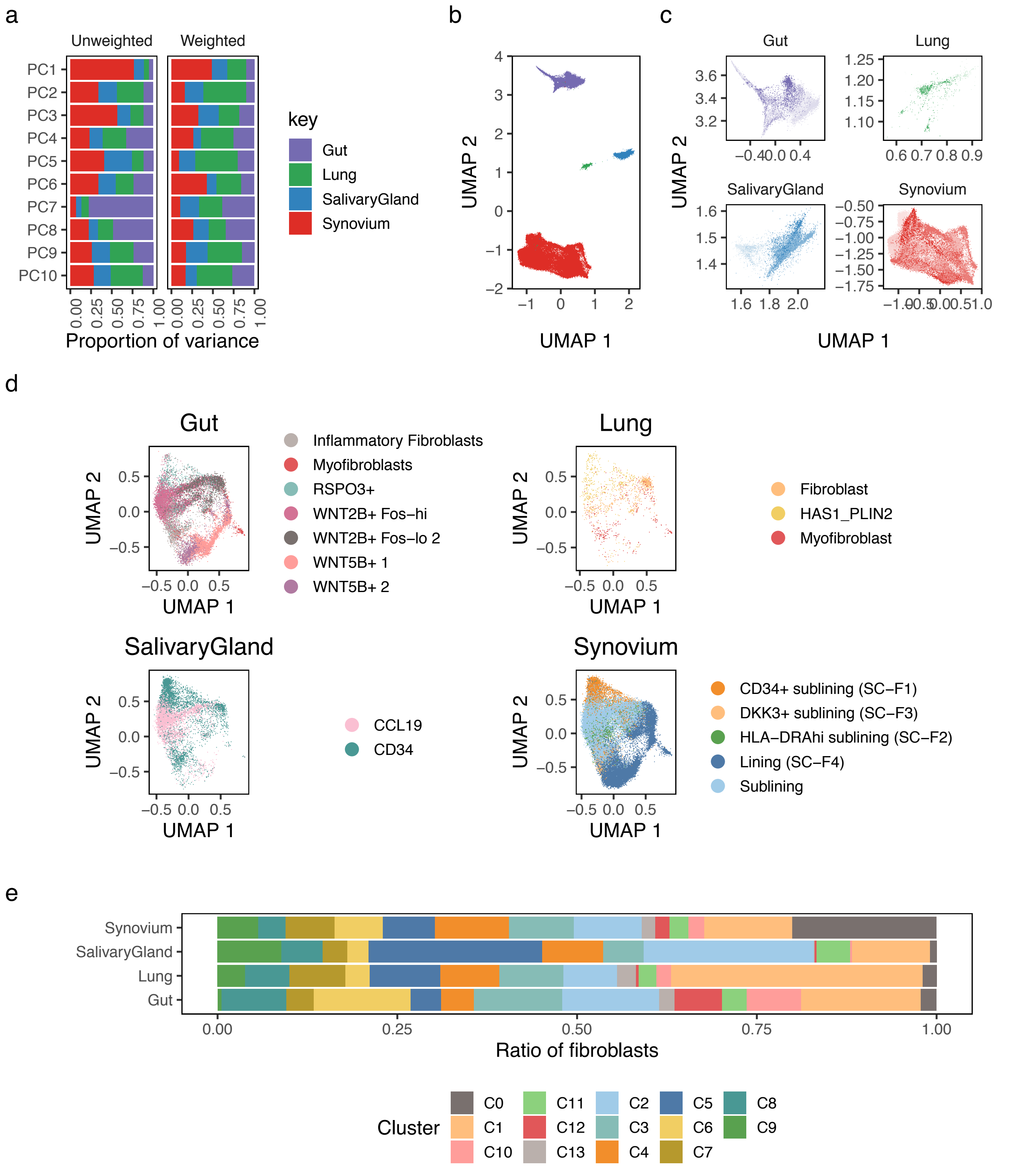


**Supplementary Figure 3. Integrated cross-tissue fibroblast reference atlas.** (a) Breakdown of variance captured in the first 10 principle components for unweighted PCA and weighted PCA shows that weighted PCA creates a more balanced embeddings among tissues. (b) Before Harmony integration, UMAP embedding of fibroblasts separates entirely by tissue. (c) Within each tissue, there is substantial separation by donor, denoted by a different hue of the corresponding tissue's color. UMAP coordinates are the same as in (b), zoomed in to focus on each tissue separately. (d) After Harmony integration, the clusters identified in tissue-specific analyses are still separated, suggesting that the Harmony embedding preserves within tissue variation. (e) Relative abundance integrative fibroblast clusters within each tissue.

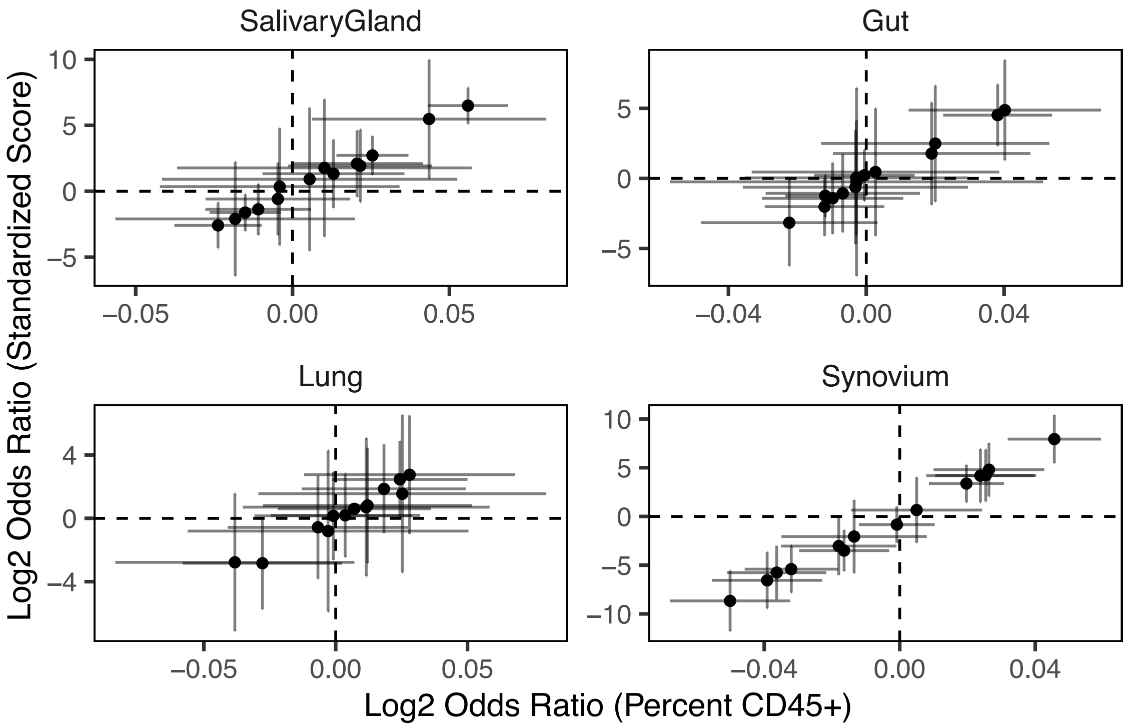


**Supplementary Figure 4. Inflammation scores.** Comparison of differential abundance analysis using raw tissue-specific scores (x-axis) and normalized cross-tissue scores (y-axis). Error bars denote 95% confidence intervals.

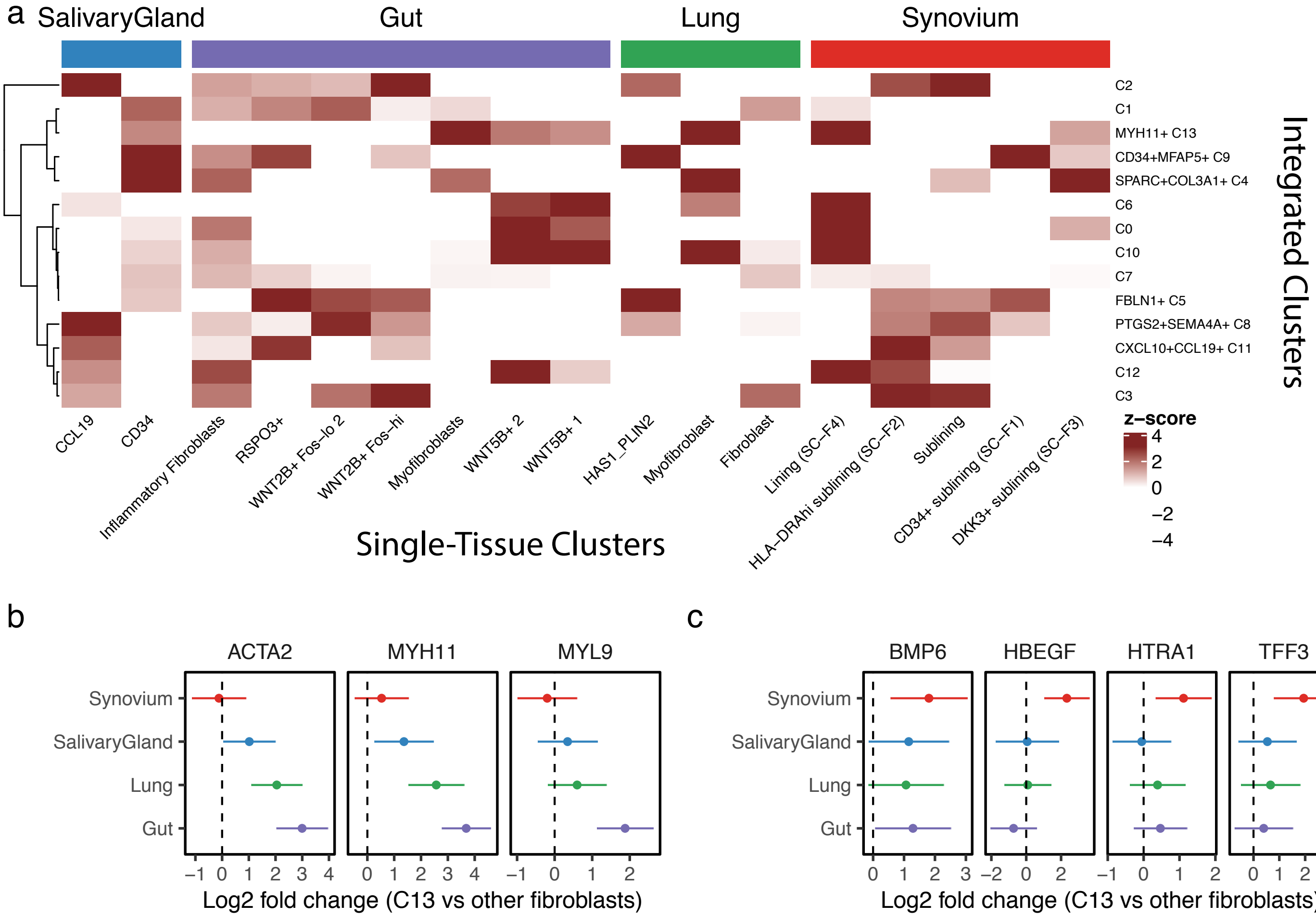


**Supplementary Figure 5. Correspondence analysis.** (a) We associated cluster identity derived in single-tissue analyses (columns) to cluster identity derived in the integrative clustering analysis (rows). Color denotes (scaled) log odds from logistic regression. (b) Gene expression fold change of genes associated with myofibroblast lineage in cluster C13 (vs other clusters). (c) Same, for genes associated with bone and cartilage repair.

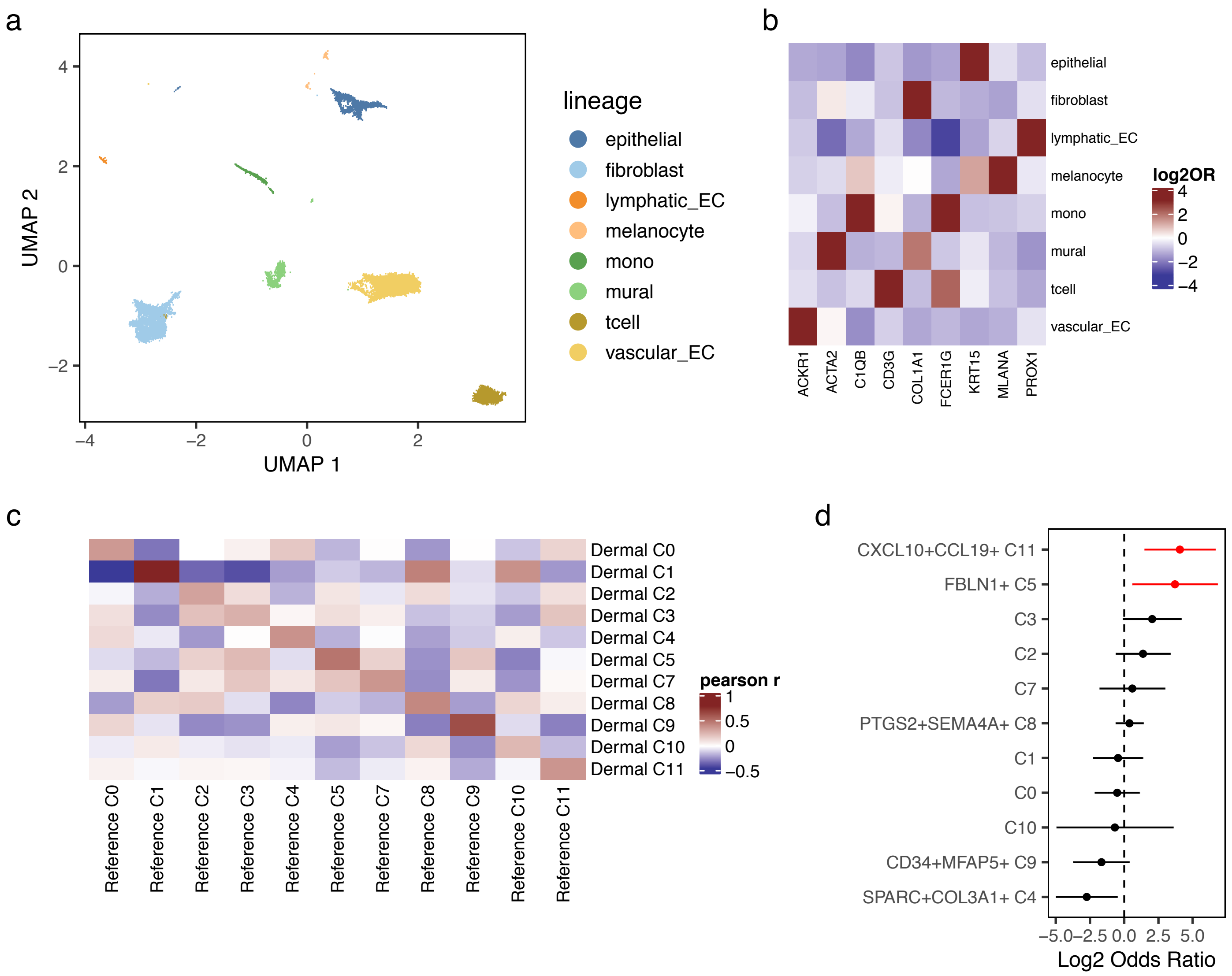


**Supplementary Figure 6. Dermal fibroblast scRNAseq profiles mapped to cross-tissue fibroblast atlas.** (a) UMAP embedding of scRNAseq profiles of skin biopsies, colored by major cell types, using (b) canonical markers: KRT15+ epithelial cells, COL1A1+ fibroblasts, PROX1+ lymphatic endothelial cells, MLANA+ melanocytes, C1QB+ myeloid cells, ACTA2+ mural cells, CD3G+ T cells, and ACKR1+ vascular endothelial cells. (c) Correlation of gene expression profiles of dermal fibroblast clusters (y-axis) against reference clusters in multi-tissue atlas (x-axis). Color denotes Pearson's correlation coefficient. (d) Differential abundance of mapped dermal fibroblast clusters with inflammation score, with 95% confidence intervals. Red denotes FDR<5%.

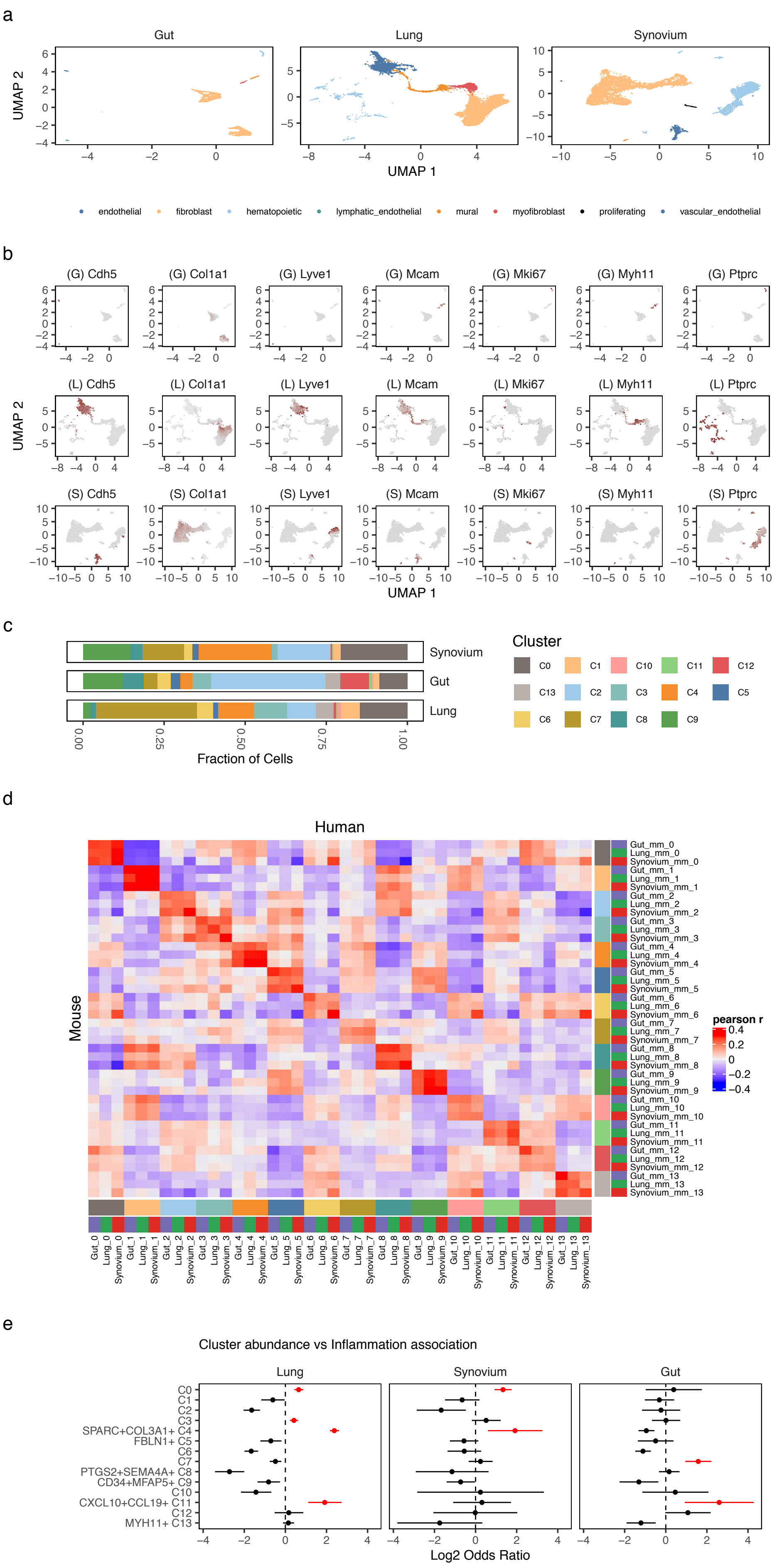


**Supplementary Figure 7. Replication in disease models.** (a) UMAP embedding of mouse scRNAseq libraries from CD45$^-$ sorted colon samples, unsorted synovial samples, and Col1a1$^+$ sorted lung samples, colored by major cell types, identified with (b) canonical markers: Cdh5+ vascular endothelial cells, Col1a1+ fibroblasts, Lyve1+ lymphatic endothelial cells, Mcam+ mural cells, Myh11+ myofibroblasts, Ki67 proliferating cells, and Ptprc+ immune cells. (c) Relative abundance of inferred fibroblast clusters in each mouse dataset. (d) Comparison of mouse cluster gene expression profiles (y-axis) to human reference cluster profiles (x-axis). Heatmap color denotes Pearson's correlation coefficient. Columns and rows are colored first by cluster identity and then by tissue. (e) Differential abundance of mapped mouse fibroblast clusters in case vs control mouse samples, with 95% confidence intervals. Red denotes FDR<5%.
